# High Content and High Throughout Phenotypic Assay for the Hourly Resolution of the Malaria Parasite Erythrocytic Cycle

**DOI:** 10.1101/2021.03.15.434938

**Authors:** Donald Bell, Sophie Ridewood, Asha P. Patel, Sun Hyeok Lee, Young-Tae Chang, Edgar Deu

**Author notes:** Corresponding author, <u>Tel:+44(0)</u> 7775 441179. These authors contributed equally to this work (Donald Bell, Edgar Deu).

## Abstract

Over the last 20 years increased funding for malaria research has resulted in very significant technical advances to study the biology of *Plasmodium* species. High throughput phenotypic assays have been developed to screen millions of compounds and identify small molecules with antiparasitic activity. At the same time, advances in malaria genetic have greatly facilitated the generation of genetically modified parasites, and whole genome genetic screens are now feasible in *Plasmodium* species. Finally, there has been an increased interest to study malaria parasites at the population level, in particular in the area of drug resistance. Drug resistant field isolates have been collected around the world, and drug resistant strains are routinely generated in the lab to study the mechanisms of drug resistance. As a result, one of the current bottlenecks in malaria research is our ability to quickly characterize the phenotype associated with compound treatment or genetic modification, or to quickly compare differences in intracellular development between strains. Here, we present a high content/high throughput phenotypic assay that combines highly selective RNA, DNA, and RBC membrane dyes to provide hourly resolution of the full erythrocytic cycle for both *P. falciparum* and *P. knowlesi*. A flow cytometry assay allows the analysis of samples in a 384-well format and a quick way to determine the parasite developmental stage. On the other hand, the fluorescence microscopy format allows for a detailed visualization of parasite morphology. Finally, using open source software we have developed protocols for the automated cluster analysis of microscopy images. This assay can be applied to any *Plasmodium* species, requires very little amount of sample, is performed with fixed cells, and is easily scalable. Overall, we believe this assay will be a great tool for the malaria community to study *Plasmodium* species.

## Introduction

Malaria is a parasitic infectious disease affecting more than 200 million people every year and killing close to half a million^1^. Malaria incidence and mortality have decreased over the last 15 years due to the distribution of insecticide-impregnated bed nets and the use of artemisinin-based combination therapies (ACTs) as the standard of care for uncomplicated malaria^2^. However, the emergence of resistance against artemisinin^3^ attests to the need to develop new antimalarials with novel mechanisms of action^4^. Encouragingly, funding for malaria research has increased over the last 20 years, which has resulted in significant scientific breakthroughs and the development of global initiatives and organizations to fight this disease.

One of these initiatives has been to partner with the pharmaceutical industry to screen millions of compounds in cell-based assays^5,6^. This approach has identified thousands of small molecules with antiparasitic activity, and follow-up studies on some of these hits has allowed the establishment of novel target- and pharmacology-based drug development programs, as well as the development of tool compounds to study parasite biology^7^. A variety of cell-based screens have been developed for *P. falciparum*^8^, and they generally fall into two categories: either they report on general parasite death or fitness by measuring metabolic activities, or they quantify parasitemia after more than 48h of treatment^9,10^. However, these assays generally do not provide information about the phenotype associated with each compound hit, thus requiring time-consuming follow-up studies. More specific microscopy-based assays have also been developed to identify compounds that block specific biological processes, such as mitochondria^11^ and apicoplast^12–14^ function, protein export^15^, or sexual^16–18^ and liver^19–23^ stage development. However, these assays often use genetically modified parasite lines and therefore are not directly applicable to any *Plasmodium* strain or species.

The recent adaptation of CRISPR/Cas9 technology^24–26^ to *Plasmodium* species, the use of novel positive and negative selection methods^27^, and the development of conditional genetic approaches^28,29^ has drastically improved our ability to quickly generate genetically modified parasites both in *P. falciparum* and *P. knowlesi*. These advances have allowed researchers to perform the first genome-wide genetic screen in *P. falciparum*^30^, and similar screens in *P. knowlesi* are likely to follow. *P. knowlesi* is a zoonotic species that is transmitted by mosquitoes from primates to humans and has recently been adapted to grow *in vitro* in human erythrocytes^31^. *P. knowlesi* is more amenable to genetic manipulation than *P. falciparum*, and because it is phylogenetically closer to other malaria-causing *Plasmodium* species (*vivax, malariae,* and *ovale*), it can be used as a surrogate for drug and vaccine development against these species^32,33^. Our increased ability to genetically modify *P. falciparum* and *P. knowlesi* allows us to generate a variety of genetically modified parasites relatively quickly, making the thorough phenotypic characterization of parasite lines one of the main bottlenecks in molecular parasitology.

In this study, we describe a novel high throughput phenotypic assay able to monitor the full erythrocytic cycle of both *P. falciparum* and *P. knowlesi* with hourly resolution. The assay combines the highly specific RNA dye 132A^34^ with Hoechst staining to simultaneously monitor intracellular parasite growth and nuclear division, respectively, as well as parasite egress and red blood cell (RBC) invasion. The samples can be analyzed either by flow cytometry or fluorescence microscopy. The flow cytometry readout allows us to run 250 samples per hour in a 384-well plate format and provides detailed information about the stage of parasite development within a 2h window. The microcopy readout allows detailed information about the morphology of the parasite. We have also used this assay to simultaneously monitor the erythrocytic cycle of multiple parasite lines in parallel in 96-well plates. Overall, this assay provides a great tool to characterize differences in development across different parasite strains and the ability to directly determine the phenotype associated with anti-parasitic compounds when performing high throughput phenotypic screens. Importantly, using the open-source software (CellProfiler and R) we have developed pipeline for the automatic analysis of microscopy images.

## Results

### Hourly temporal resolution of the *P. falciparum* erythrocytic cycle by flow cytometry

A culture of 3D7 *P. falciparum* was tightly synchronized within a 1h window and grown for 80h in round-bottom 96-well plates. Aliquots were collected at different time points between 1 and 80 hours post invasion (hpi), fixed for 1h, diluted 10-fold in PBS, and stored at 4°C until further analysis. Samples were then diluted 10-fold in PBS in round bottom 384-well plates and stained simultaneously with 1 μg/mL Hoechst, and 2 μM 132A for 30 min at 37°C. Ten microliters of sample were then run in a Fortessa X20 (BD) equipped with a plate autosampler. Hoechst staining was measured using a UV (355 nm) laser and a 450/50 nm emission filter, and 132A staining with a blue (488 nm) laser and a 610/20 nm emission filter.

Fig. 1A shows representative flow cytometry plots at selected time points. The large cell population with low DNA and RNA signals corresponds to uninfected RBCs (uRBCs). Newly infected RBCs (iRBCs) run as a population positive for DNA staining and with an RNA signal that is about two-fold higher than the one observed for uRBCs. As the parasite undergoes intracellular development the population of iRBCs initially increases its RNA signal with little change in DNA levels, i.e. transition from ring to trophozoite stages between 1 and 30 hpi. This is then followed by a rapid exponential increase in both DNA and RNA signals illustrating nuclear division and continuous parasite growth within iRBC during schizogony.

**Figure 1.**
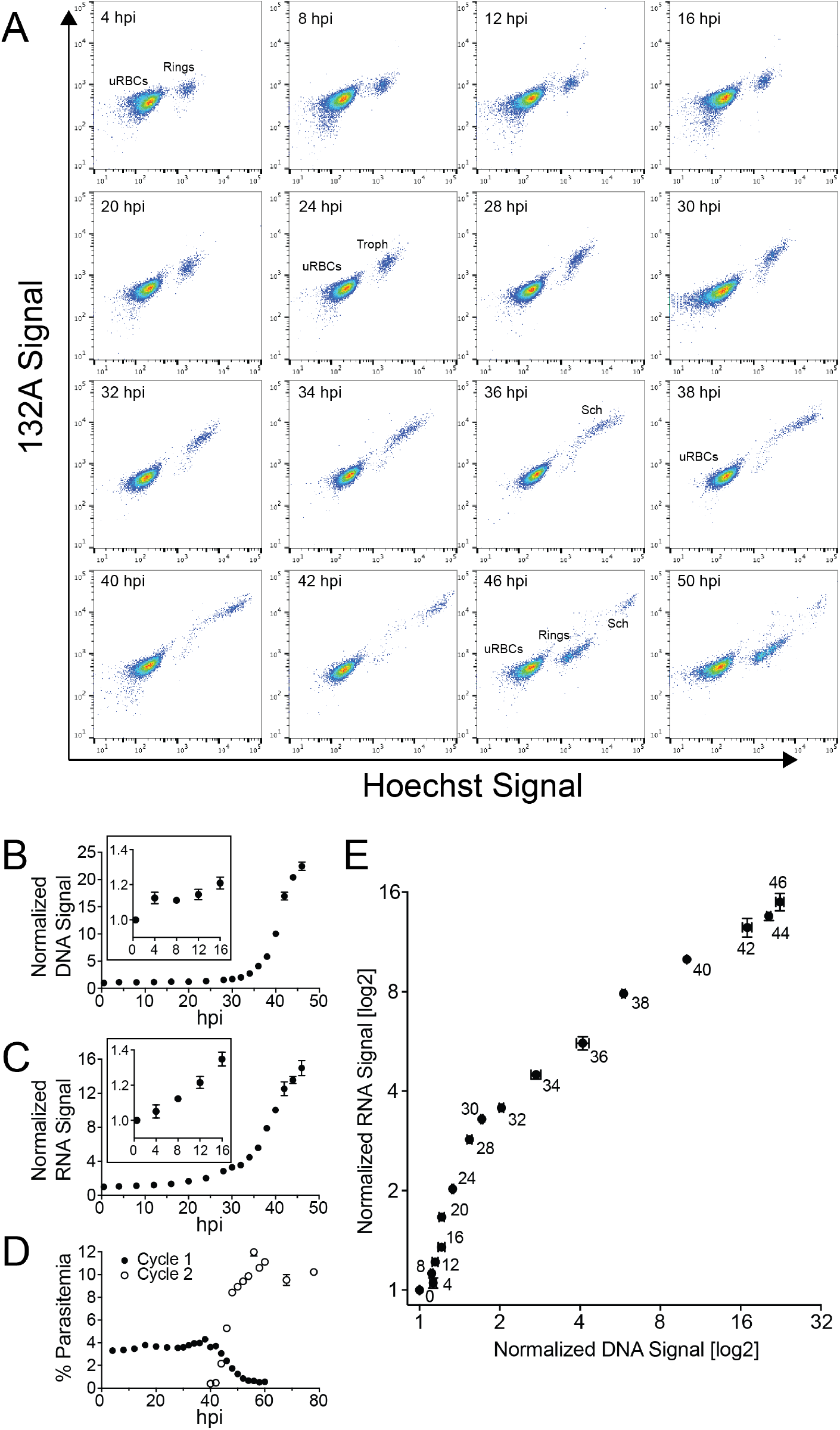
Flow cytometry analysis of *P. falciparum* erythrocytic cycle. A tightly synchronized culture of 3D7 *P. falciparum* parasite at 3% parasitemia was culture at 37°C in 96-well plates for 80h. Samples were collected at different time points between 0 and 80 hpi, fixed, stained with Hoechst and the RNA dye 132A, and analyzed by flow cytometry. (**A**) Raw data flow cytometry plots showing the different populations of infected and uninfected RBCs at different time points (indicated in the upper left of each graph). X- and Y-axes show the fluorescence signal in the Hoechst and 132A channels, respectively. The cell population on the lower left corresponds to uRBCs and those positive for DNA staining to iRBCs. Populations of iRBCs corresponding to ring, trophozoite and schizont stage parasites are indicated. (**B-C**) The time-dependence of the median fluorescence intensity normalized to the 1 hpi time point for the DNA and RNA signals are shown in panels **B** and **C**, respectively. Insets illustrate changes in signal over the first 16h of the erythrocytic cycle. (**D**) Quantification of parasitemia, egress and invasion. Because of the very significant difference in DNA and RNA signals between rings and schizonts, we were able to differentiate parasites before egress (Cycle 1, filled circles) and after invasion (Cycle 2, empty circles). This allowed us to measure egress (decrease in the schizont population) and invasion (increase of the ring population) over time and determine parasitemia at each cycle. (**E**) Graph showing how the DNA (X-axis) and RNA (Y-axis) signals varied overtime (hpi are shown next to each data point) throughout intracellular parasite development (0-48 hpi). For **B-D**, error bars present standard error of three technical replicates.

The time-dependence of the DNA and RNA median signals of iRBCs, normalized to the initial time point (1 hpi), are shown in Fig. 1B and C, respectively. While there are no significant changes in the DNA signal within the first 24h of the cycle, significant differences in RNA levels could be quantified within the first 8h. We think that the small increase in DNA signal observed between 12 and 28 hpi is likely due to parasites becoming metabolically more active at trophozoite stage. Increase in transcription levels requires a decrease of chromatin condensation that likely increases Hoechst binding to DNA.

Note that the difference between mature schizonts and newly iRBCs is very pronounced (see 46 hpi plot in Fig. 1A) allowing us to monitor egress and invasion. As shown in Fig. 1D, the decrease in schizont population, i.e. egress, occurs between 42 and 54 hpi, and it is concomitant with the increase in the number of ring stage parasites, i.e. invasion. The graph in Fig. 1E shows changes in median DNA vs RNA levels as a function of time and reflects the path followed by the iRBC population across the flow cytometry plot, i.e. initial increase in RNA levels between 1 and 30 hpi, followed by an increase in DNA and RNA levels during schizogony.

### Analysis of the *P. falciparum* erythrocytic cycle by fluorescence microscopy

We then used the same samples to monitor the full parasite life cycle by microscopy. In addition to Hoechst and 132A staining, we used wheat germ agglutinin conjugated to Alexa 647 (WGA-647) to label the surface of the RBC. This provides better contrast to delineate the RBC membrane than bright field images and allowed us to use imaging software for automatic recognition of the RBC surface. Fig. 2 shows representative images obtained across the *P. falciparum* erythrocytic cycle. A clear increase in the size and intensity of the RNA staining is observed throughout the erythrocytic cycle, illustrating continuous parasite growth within iRBCs. Note that the hemozoin crystal can be detected on the bright field images from 24 hpi onwards, and that the circular depression observed at ring stages^35^, which does not colocalize with the nucleus or the food vacuole, is clearly visible at the 12 and 16 hpi time points. Between 32 and 46 hpi, a clear increase in the number of nuclei per iRBC is observed, thus reflecting nuclear division during schizogony. Note that in the image shown at 44 hpi, the RNA signal is continuous and surrounds each of the nuclei within the iRBC. However, the one shown at 46 hpi shows RNA signal enclosed within each mature merozoites. Therefore, this assay could be used to monitor cytokinesis in *P. falciparum*. Finally, during egress and invasion we observed a lot of newly iRBCs showing the typical amoeboid physiology. Although this is observable in the bright field, the morphology of the parasite is more clearly observable in the RNA channel (See representative image at 48 hpi).

**Figure 2.**
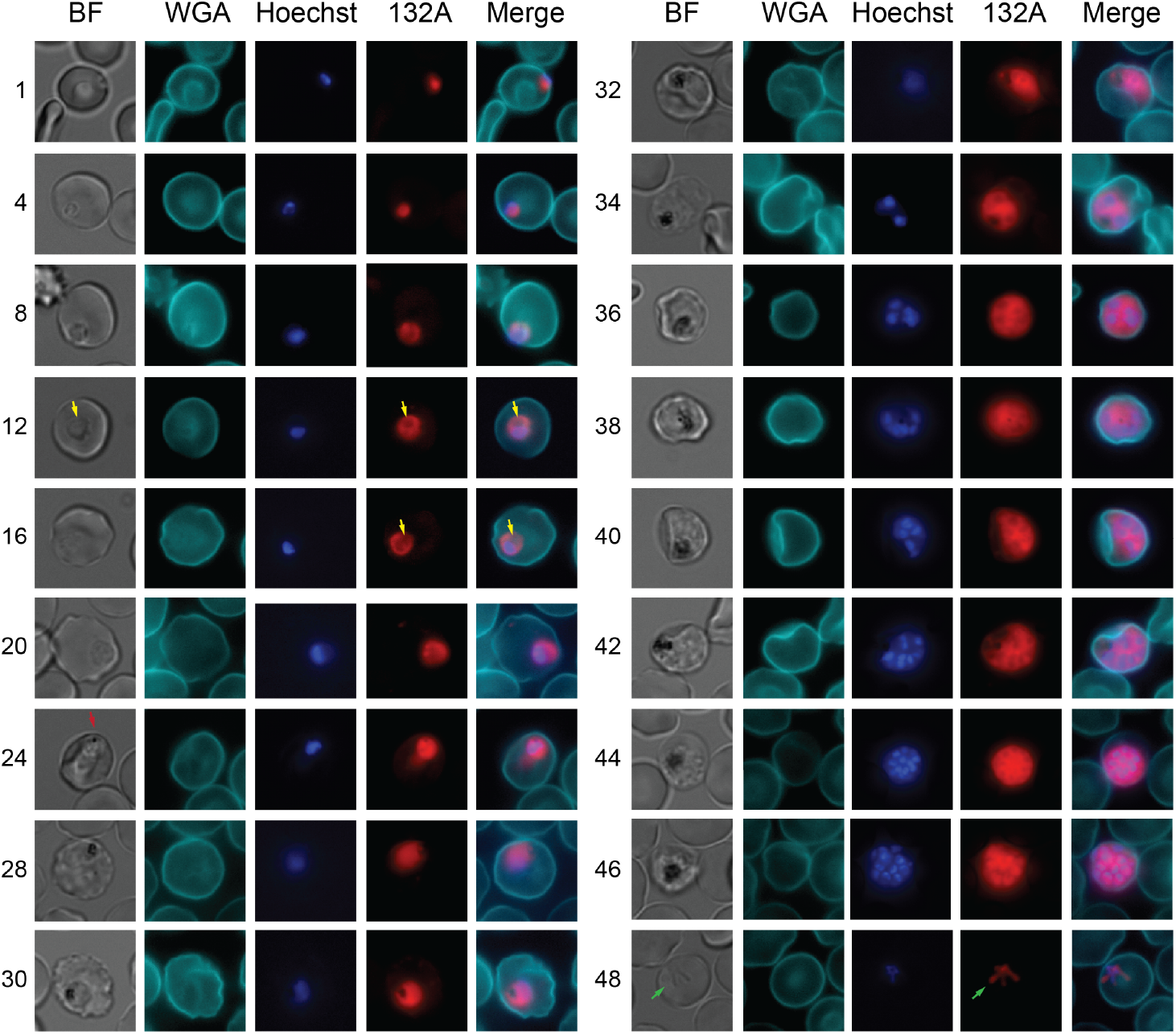
Microscopy analysis of *P. falciparum* erythrocytic cycle. Fixed samples of *P. falciparum* 3D7 parasites were stained with Hoechst, the RNA dye 132A, and WGA-647. The latter binds to lectins on the surface of the RBC and allows us to clearly visualize the RBC membrane. Cells were allowed to settle at the bottom of a glass bottom 96-well plate and were imaged in an inverted microscope. Representative images obtained using the bright field (BF), WGA-647 (WGA), Hoechst, and 132A channels are shown at each of the indicated time points, as well as a merge images of the fluorescent channels. The yellow arrows at 12 and 16 hpi point to the circular cavity observed in ring stage parasites. The red arrow at 24 hpi indicates the first time point at which we consistently observed hemozoin formation in the BF images. Note that in the 46 hpi image fully formed merozoites can clearly be observed as distinct circular RNA staining around each of the nuclei. The green arrows at 48 hpi point to newly invaded RBCs with a characteristic amoeba shape rather than a spherical morphology (1hpi).

### Evaluation of the assay

Because of the very tight temporal resolution of this assay, we systematically use it to characterize genetically modified parasite lines in the lab. In particular, our lab has been using the DiCre system to conditionally knock out different genes of interest in *P. falciparum*. When we set up this time course, in addition to the 3D7 line, we included in our experiment the 1G5 line that endogenously expressed DiCre^36^, as well as the previously characterized DPAP3cKO line^37^. DPAP3 is a cysteine protease that is important for efficient RBC invasion. Our previous results showed that conditional KO of DPAP3 has no effect in intracellular parasite development or egress but results in a 50% decrease in RBC invasion^37^. We used these three lines to evaluate the consistency of our assay across different cell lines, and to determine whether rapamycin (RAP) treatment has any effect in parasite development.

Each line was treated for 3h with DMSO or 100 nM RAP at 1 hpi, and an aliquot of each culture was taken at different time points and analyzed by flow cytometry as described above. No significant differences in intracellular development, measured by RNA or DNA levels, nor in the time of egress were observed when comparing DMSO vs. RAP treatment for these three lines. As expected, we observed about a 50% decrease in invasion efficiently for the DPAP3cKO line upon RAP treatment. Note that we observed a consistent 10% drop in invasion efficiency upon RAP treatment on the 3D7 and 1G5 lines. Since no significant differences in intracellular parasite development were observed between these 3 lines in the presence or absence of RAP, we performed t-test analysis of these six time courses to determine whether the level of DNA and RNA signals were significantly different between consecutive time points (Fig. 3A). For almost all consecutive time points between 0 and 44 hpi we measured a significant increase in RNA level. On the other hand, increase in DNA levels was only significant between 20 and 44 hpi. As mentioned above, we think that the increase in Hoechst staining between 20 and 32 hpi is likely due to a decrease in chromatin condensation allowing better DNA staining since our microscopy images show that iRBC only have one nucleus at these time points. Interestingly, we observed a small but significant decrease in RNA levels between 46 and 48 hpi, which might be due to an overall decrease in transcription in mature merozoites immediately before parasite egress. Overall, the comparison of the time course assay across different cell lines attests to the consistent and high temporal resolution of this assay.

**Figure 3.**
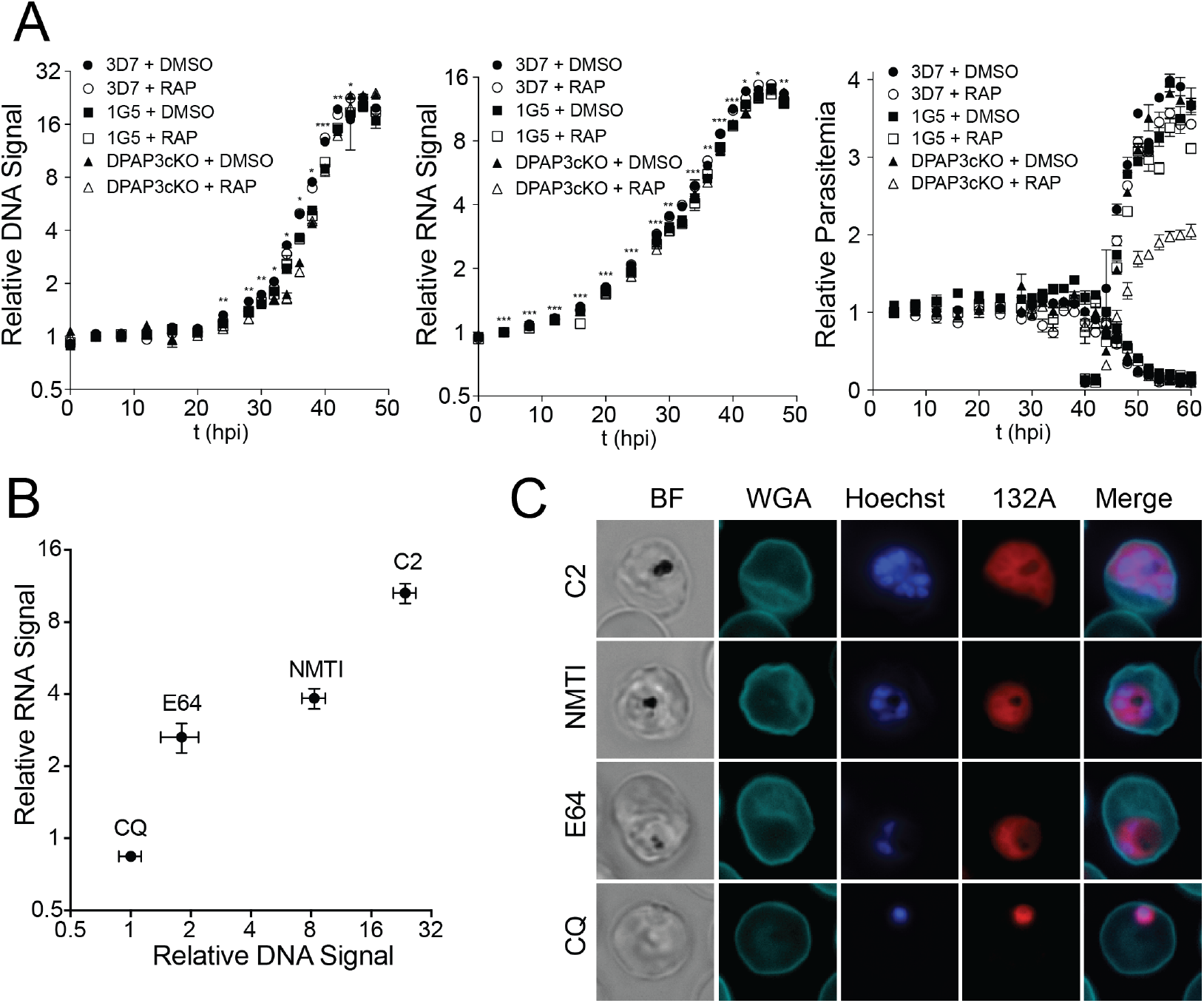
Time course reproducibility and antiparasitic compounds phenotype. (**A**) Cultures of tightly synchronized 3D7, 1G5, or DPAP3cKO parasites were treated between 1 and 4 hpi with DMSO or RAP and cultured for 80h in 96-well plates in triplicate. Aliquots of each culture were collected at different time points, fixed, stained with Hoechst and 132A, and analyzed by flow cytometry. The time-dependence change in DNA, RNA and parasitemia are shown in the left, middle and right graphs respectively. Asterisks indicate t-test significant values of the average changes in DNA or RNA levels between consecutive time points. (**B**) Synchronized cultures of 3D7 parasites were treated with CQ, E64 or an NMTI at ring stage parasites and with C2 at schizont stage. Cultures were fixed at 48 hpi, stained with Hoechst and 132A, and analyzed by flow cytometry. The median DNA vs RNA signal is shown in the graph for each treatment. Error bars correspond to standard error of nine replicates. (**C**) Representative fluorescence microscopy images showing the phenotype associated with CQ, E64, NMTI or C2 treatment: pyknotic arrest at ring stage, swollen food vacuole at trophozoite stage, arrest at mid schizogony, or block of parasite egress, respectively.

One of the potential applications of this assay is to perform high throughput phenotypic screens. The flow cytometry-based assay can be performed in 384-well plates allowing running around 250 samples per hour. The flow cytometry readout provides a rapid way to identify compounds that significantly arrest or delay parasite development, as well as information about the developmental stage affected. To get a better evaluation of the phenotype associated with each hit compound, the same samples can then be imaged by microscopy. To test whether our assay could discriminate and quantify different phenotypes, a synchronous culture of 3D7 parasite was treated with four anti-parasitic compounds known to arrest development at different stages. Chloroquine (CQ) quickly kills parasites at ring stage. E64 is a general cysteine protease inhibitor that inhibits the falcipains and delays parasite development at trophozoite stage. Inhibition of the falcipains blocks the haemoglobin degradation pathway resulting in an engorgement of the food vacuole due to the accumulation of undigested haemoglobin in this organelle^38^. Treatment of ring stage parasites with the N-myristoyl transferase inhibitor (NMTI) has been shown to arrest parasite at mid schizogony^39^. Finally, the cGMP-dependent protein kinase inhibitor Compound 2 (C2) blocks egress of very mature segmented schizonts^40^. Figure 3B shows the median RNA and DNA level obtained from our FACS analysis and illustrate how each compound arrests development at a different stage along the flow cytometry path depicted in Fig. 1E. We then looked at these arrested parasites by microscopy and clearly were able to observe the expected phenotypes (Fig. 3C): pyknotic morphology for CQ treatment, trophozoites containing a swollen food vacuole with E64, schizonts containing 5-7 nuclei for parasites treated with the NMTI, and fully segmented schizonts arrested before egress with C2.

### Analysis of the *P. knowlesi* erythrocytic cycle by flow cytometry and fluorescence microscopy

We then evaluated the ability of our assay to monitor the *P. knowlesi* erythrocytic cycle. *P knowlesi* parasites are harder to synchronize because ring-stage parasites are not resistant to sorbitol treatment, and schizont removal using Percoll gradient is not as efficient as in *P. falciparum*. In order to obtain a tightly synchronized *P. knowlesi* culture, schizont stage parasites were first purified using a Percoll gradient and cultured with RBC for 4h to allow the most mature population of parasites to egress and reinvade. A second schizont purification was then performed, and these schizonts were incubated for 4h with 1 μM of C2 to obtain a population of highly mature schizonts. After C2 washout out, schizonts were incubated with RBCs for 2h under shaking condition to maximize egress and invasion. The culture of newly iRBCs was then collected using another Percoll gradient. By using this method we were able to obtain a culture enriched with early ring stage parasites at 5% parasitemia. However, this culture still contained a small proportion of schizonts (around 3.5% of iRBCs, see 1 hpi flow cytometry plot in Fig. 4A). Aliquots from this culture were then collected every 1 to 2h between 1 and 34 hpi.

**Figure 4.**
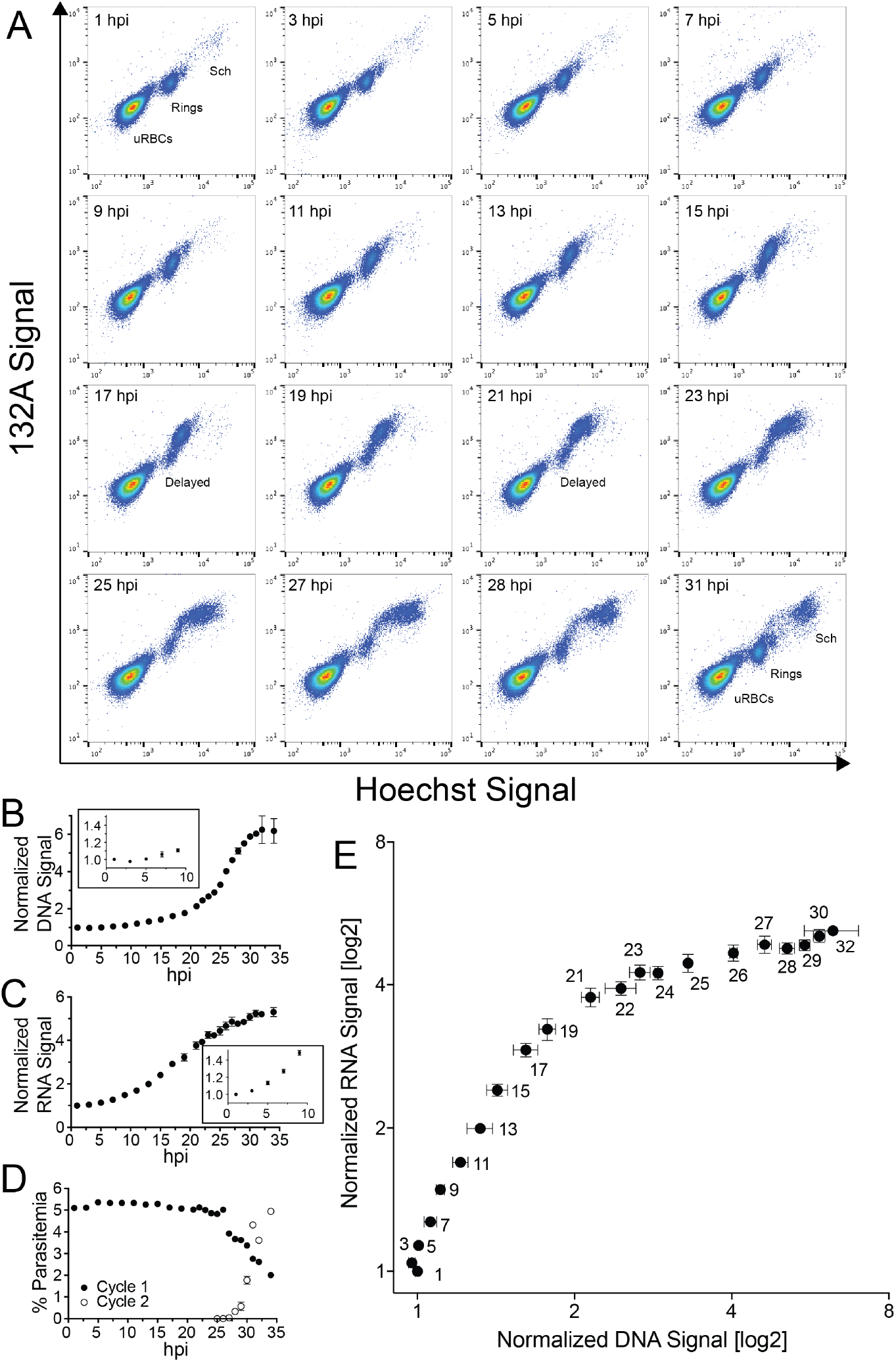
Flow cytometry analysis of *P. knowlesi* erythrocytic cycle. A tightly synchronized culture of A1-H.1 *P. knowlesi* parasites at 5% parasitemia was cultured at 37°C for 35h. Samples were collected at different time points, fixed, stained with Hoechst and 132A, and analyzed by flow cytometry. (**A**) Raw data flow cytometry plots showing the different populations of infected and uninfected RBCs at different time points (indicated in the upper left of each graph). X- and Y-axes show the fluorescence signal in the Hoechst and 132A channels, respectively. The cell population on the lower left corresponds to uRBCs and those positive for DNA staining to iRBCs. Populations of iRBCs corresponding to ring, trophozoite and schizont stages are indicated. A population of delayed parasites is clearly observable between 17 and 25 hpi and corresponds to newly iRBCs from the small population of schizonts that was present at 1 hpi. (**B-C**) The time-dependence of the median fluorescence intensity normalized to the 1 hpi time point for the DNA and RNA signal are shown in **B** and **C**, respectively. The insets illustrate the change in signal over the first 9h of the erythrocytic cycle. (**D**) Quantification of parasitemia, egress and invasion. Because of the very significant difference in DNA and RNA signal between rings and schizonts, we were able to differentiate parasites before egress (Cycle 1, filled circles) and after invasion (Cycle 2, empty circles). This allowed us to measure egress (decrease in the schizont population) and invasion (increase of the ring population) over time, and quantify changes in parasitemia at each cycle. (**E**) Graph showing how the DNA (X-axis) and RNA (Y-axis) signals varied over time (hpi are shown next to each data point) throughout intracellular parasite development (0-32 hpi). For **B-D**, error bars present standard error of three technical replicates.

Raw data flow cytometry plots at selected time points are shown in Fig. 4A. Note that the small population of schizonts present at 1 hpi disappears between 3 and 9 hpi, reflecting parasite egress and RBC invasion. From 15 hpi we observed a small distinct population of delayed parasites that displayed lower DNA and RNA levels than the main iRBCs population. We think that this delayed population corresponds to RBC that were infected as a result of the small population of schizonts present at 1 hpi. This delayed population of iRBCs was excluded from our analysis whenever it could be differentiated from the main population, i.e. between 15 and 23 hpi. Figure 4B and C show the time-dependence of DNA and RNA levels in *P. knowlesi* iRBCs, respectively. As for *P. falciparum*, only the RNA dye allows to monitor early parasite development at ring stage. Fig. 4D shows that under our culturing conditions the time of egress and invasion occurs around 32 hpi. Finally, Fig. 4E shows the path the parasites take over the DNA *vs.* RNA flow cytometry plot over time, which illustrates significant differences with the one measured for *P. falciparum* (Fig. 1E). Similarly to *P. falciparum*, we measured a 2-fold increase in Hoechst signal over the first half of the cycle probably due to increase transcription at trophozoite stage. This is followed by an exponential increase in DNA levels during schizogony. However, while RNA levels increased exponentially during schizogony in *P. falciparum* (30-46 hpi), in *P. knowlesi* RNA levels increase linearly over the equivalent period (20-32 hpi). These differences might reflect the fact that *P. knowlesi* produces less merozoites per iRBC than *P. falciparum*, and therefore, it might not require sustained parasite growth throughout the erythrocytic cycle. Note that the maximum level of DNA signal obtained for *P. knowlesi* is only 6-fold higher than that of newly iRBC as opposed to 22-fold for *P. falciparum*, thus reflecting the number of merozoites produced per schizont for each of these species.

The same samples used to monitor the full life cycle of *P. knowlesi* by flow cytometry were stained with Hoechst, 132A and WGA-647 for fluorescence microscopy analysis. Representative images at different hpi are shown in Fig. 5. As for *P. falciparum*, the indentation at ring stage is observable in the RNA channel at 11 hpi. Consistent with the flow cytometry results, we did not observe a big increase in the area or intensity of the RNA signal during schizogony as was observed in *P. falciparum.* Fully segmented schizonts can be observed at 29 hpi in the RNA channel indicating invagination of the plasma membrane to form individual merozoites. At 27 hpi the RNA signal shows a continuous circular pattern surrounding multiple nuclei, consistent with nuclear division before cytokinesis. Interestingly, at 28 hpi we could observe star-shape RNA signal, which we believe correspond to schizonts undergoing cytokinesis.

**Figure 5.**
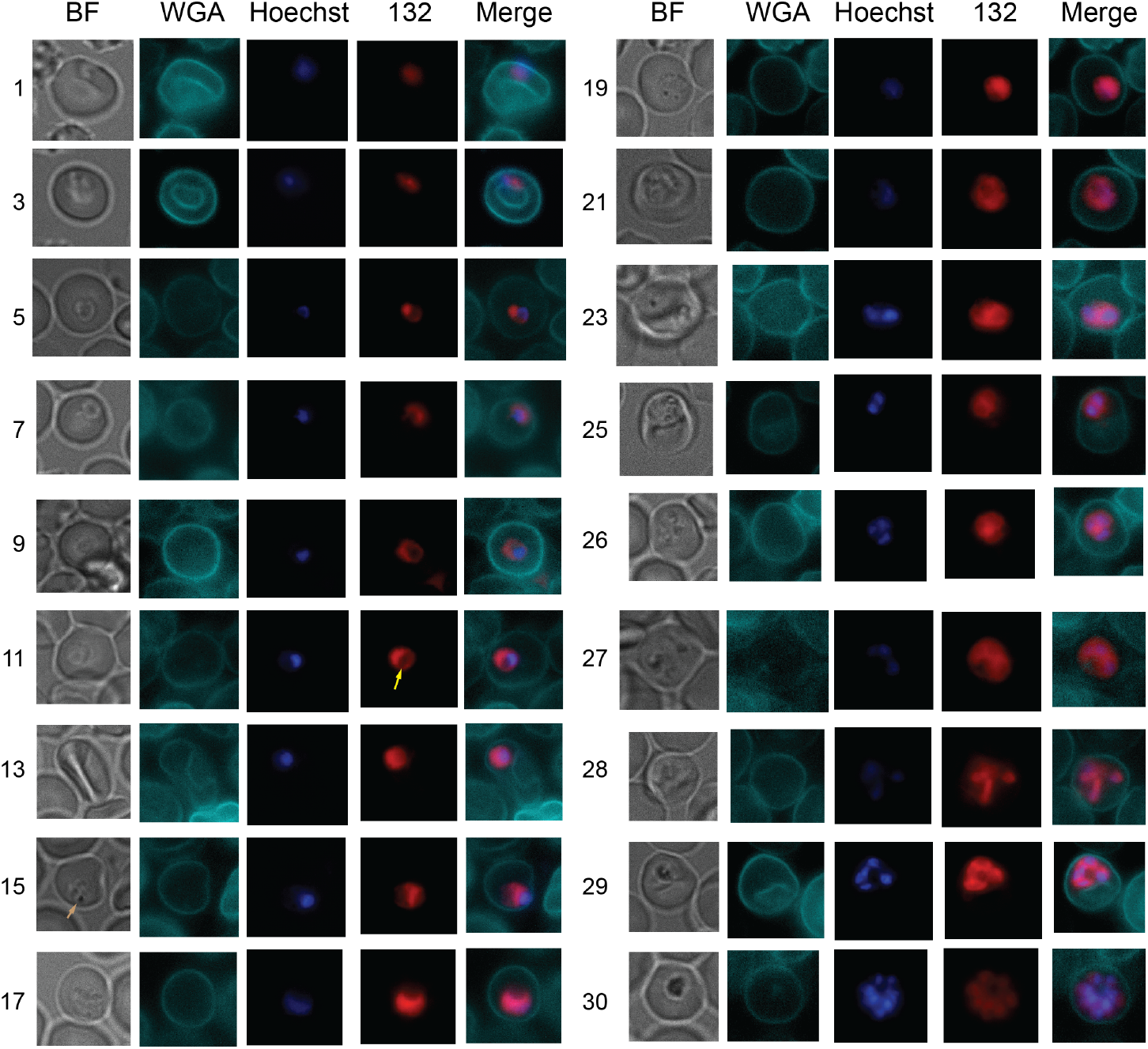
Microscopy analysis of *P. knowlesi* erythrocytic cycle. *P. knowlesi* fixed samples were stained with Hoechst, 132A, and WGA-647. Cells were allowed to settle at the bottom of a #1.5 glass bottomed 96-well plate and were imaged in an inverted microscope. Representative images obtained using the bright field (BF), WGA-647 (WGA), Hoechst, and 132A channels are shown at each of the indicated time points. Merge images of the fluorescent channels are also shown. The yellow arrow at 11 hpi points to the circular cavity observed in ring stage parasites. The brown arrow at 15 hpi indicates the first time point at which we consistently observed hemozoin formation in the brightfield. Note that in the 29 hpi image, fully formed merozoites can clearly be observed as distinct circular RNA staining surrounding each of the nuclei.

### Automated cluster analysis of *P. knowlesi* images

Labeling of RBC with WGA-647 allowed us to achieve high contrast to delineate the RBC membrane. We used CellProfiler to identify every single RBC in the field and generate a cropped image of each iRBC, defined as RBCs containing DNA and RNA signal. At this stage CellProfiler was used to quantify parasitemia and the number of nuclei per iRBC, and to generate a database containing a variety of image analysis parameters for each of the segmented iRBCs, such as area, shape, intensity and texture in each of the microscopy channels, i.e. bright field, DNA, RNA, and WGA-647. We then used R software to automatically classify the iRBCs into 12 different clusters through multiple component analysis (Fig. 6A). Images in each cluster were visually inspected to exclude problematic images for further analysis, i.e. images showing multiply infected RBCs or of poor quality (out of focus, lacking signal in one or more channels, or containing more than one RBC). We noted that cluster XII had only 20 images, and most of them correspond to problematic images. We therefore disregarded cluster XII in most of our analysis. For the purpose of this paper, the clusters were numbered according to the stage of intracellular development from early rings (I) to mature schizonts (XI). As can be seen in Fig. 6A, the cluster follows an elliptical pattern in the 3D t-SNE graph and their position seems to be consistent with the parasite developmental stage.

**Figure 6.**
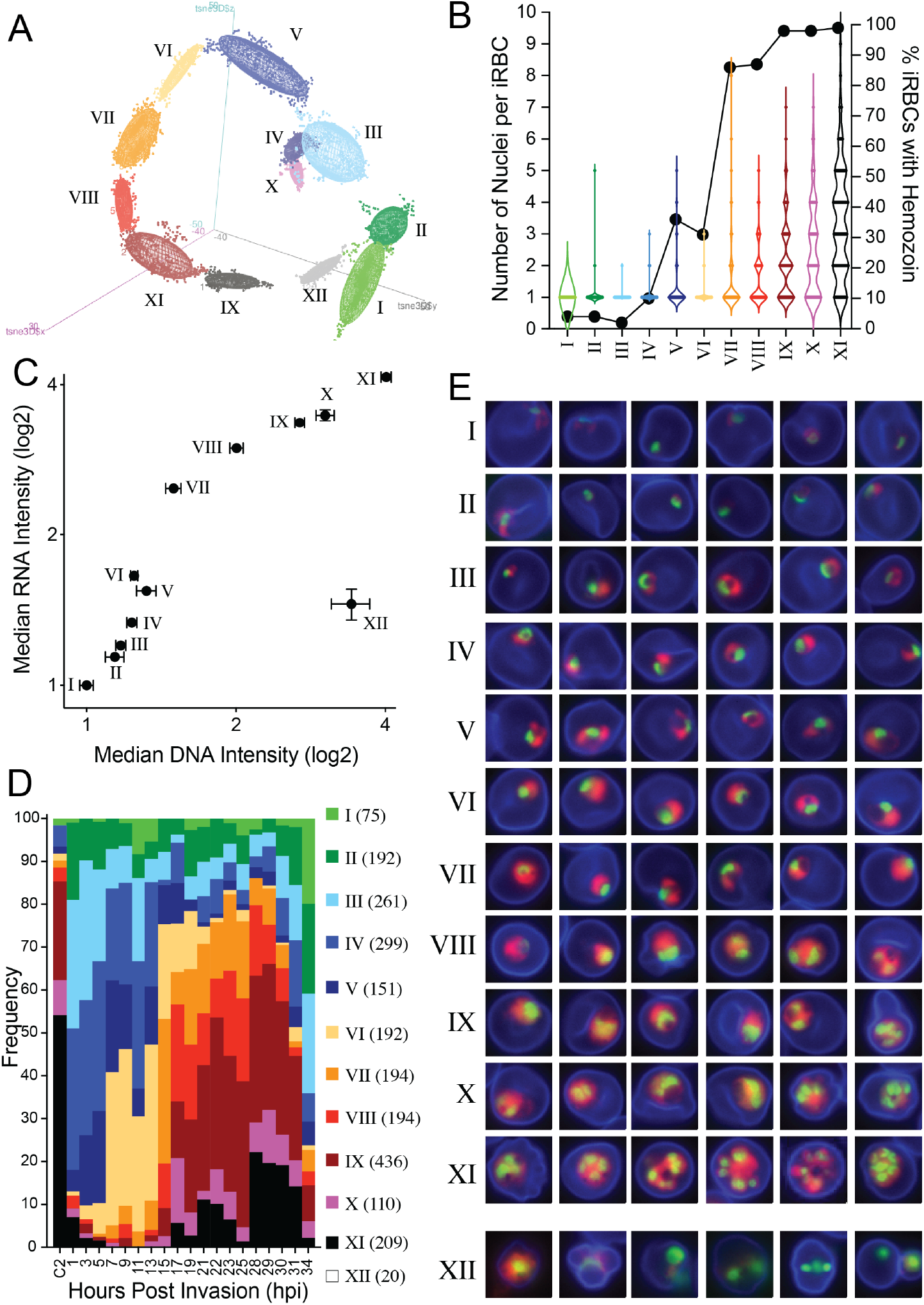
Automated cluster analysis of *P. knowlesi* intracellular development. **(A)** We used the statistical computing software R to classify the cropped images of iRBCs obtained for the *P. knowlesi* time course in 12 different clusters based on the images obtained in four different channels: brightfield, WGA-647, Hoechst and 132A. The 3D graph shows the distribution of the clusters based on 3 component analysis (t-SNE). (**B**) For each cluster, the number of nuclei in each iRBC and whether or not it contained hemozoin crystals was quantified visually. (**C**) The median DNA intensity of each iRBC image in each cluster was plotted versus the median RNA intensity. Consistent with our flow cytometry results, we observed an increase in RNA intensity for the clusters belonging to the early stages of parasite development (rings and trophozoites), followed by an increase in both DNA and RNA levels during schizogony. Note that cluster XII is an outlier in this plot because most of the images in this cluster were problematic. (**D**) For each time point, the proportion of iRBCs belonging to each cluster (I-XI) was quantified and normalized to 100%. Note that the first bar in the graph corresponds to C2 arrested parasite, i.e. very mature schizonts. The number of images in each cluster is shown in parenthesis in the legend. (**E**) The representative images for each cluster are shown as merged images across the three fluorescence channels (WGA-647 in blue, Hoechst in green and 132A in red).

To control that indeed our automated cluster analysis was classifying iRBCs images according to parasite development, we counted the number of nuclei in each iRBCs and the proportion of iRBCs where we could observe hemozoin crystal(s) in the BF channel (Fig. 6B). Infected RBCs in clusters I to VII contain mainly a single nucleus, and therefore likely correspond to the ring to trophozoite transition. From cluster VIII to XI we observed an increased in the number of nuclei per iRBCs, illustrating schizogony. Hemozoin crystals are practically absent in the iRBC images of clusters I through III, and the proportions of iRBCs having hemozoin gradually increase from cluster IV through VIII, indicating trophozoite stage. As expected, close to 100% of iRBCs have hemozoin crystals in clusters IX to XI, which contain mainly schizont stage images. We then plotted the median DNA intensity obtained for each iRBC image as a function of median RNA intensity for each cluster (Fig. 6C). This 2D graph is very similar to the one obtained using our flow cytometry analysis (Fig. 4E) and indicates that clusters I to VII correspond to the early ring to late trophozoite transition, and VIII to XI to schizogony. Importantly, the distribution of each cluster at each time point is consistent with the expected distribution of the different parasite stages at different hpi (Fig. 6D).

Representative images of iRBCs in each cluster are shown in Fig. 6E. Clusters I-IV correspond to ring stage parasites. In the first two clusters nuclear staining is generally more prominent than RNA staining and occupies more space in the iRBCs. In cluster I, a significant number of ring stage parasites show an amoeboid structure (25%), indicating that this cluster contains images of newly infected RBCs. In cluster III and IV, the RNA signal becomes more intense and larger, and the ring shape of the iRBCs, with the circular indentation, is clearly observable in 25% of the images. Images in clusters V, VI, and VII correspond to the late ring to trophozoite transition, characterized by an increase in RNA staining and the increase presence of hemozoin crystals (Fig. 6B). The main difference between cluster V and VI is in the pattern of RNA staining. In cluster V, RNA staining is quite irregular, patchy and surrounding the nuclei. In cluster VI, most images display RNA staining in one side of the image and mainly as a single spherical stain. From cluster I through VII, a single nucleus is generally present in iRBCs. Starting at cluster VIII and until cluster XI, we observed an increase in the number of nuclei and/or a significant increase in the size of nuclear staining, either representing dividing nuclei or overlapping nuclei. In C2-treated parasites that were arrested before egress, 54% of the images were classified in cluster XI (Fig. 6D). In this cluster, 42% of images correspond to segmented schizonts, as opposed to only 11% in cluster X. Overall, these results illustrate that automated and unbiased cluster analysis of the images obtained using this assay is consistent with parasite development and could be adapted to identify specific phenotypes of interest.

## Discussion

The high content phenotypic assay presented here allows unprecedented hourly dissection of the erythrocytic cycle of both *P. falciparum* and *P. knowlesi* by either flow cytometry or fluorescence microscopy. One of the advantages of this assay compared to other phenotypic assays is that it is performed in fixed cells rather than in live parasites. This makes this assay easier to implement since it does not require having microscopes or cytometers in a containment facility. Also, because fixed samples are stable at 4°C for months, it allows researchers to collect large amounts of samples without the time constraints imposed by live microscopy assays. In addition, our fixation and staining protocol does not require any washes, making the handling of samples and plates straightforward and readily scalable. Finally, the fact that this assay uses fluorescent dyes makes it broadly applicable to any parasite line and species, as opposed to other type of microscopy-based phenotypic assays that require genetically modified cell lines with fluorescent protein reporters.

When using the flow cytometry readout using a 384-well plate set-up, we can run about 250 samples per hour. Although this assay is probably not amenable for large high throughput screening campaigns of millions of compounds, it is suitable for the screening of several thousands. The microscopy-based readout is more time consuming and therefore probably not suitable for HTS. However, it is an excellent assay to validate hits identified by flow cytometry or to characterize different parasite cell lines in parallel in a very efficient way. Another potential application of this flow cytometry-based assay could be to monitor the development of a variety of parasite strains in response to drug treatment. This can be especially interesting when comparing the response of drug-resistant strains, either field isolates or lab generated strains, to increasing concentrations of drugs.

We believe our method to perform automatic image analysis of iRBCs will be of great utility to the malaria research community. This method was developed exclusively using open-source software so that it can be implemented right away in any lab using the codes provided in the supplementary materials. As illustrated from our cluster analysis, this unbiased method allows clear differentiation of different stages of parasite development. Importantly, similar protocols either using CellProfiler Analyst or the unbiased cluster analysis can be adjusted to reflect the different types of expected phenotypes in any assay. For example, our images of iRBCs presented in this manuscript clearly allow us to differentiate different phenotypes during parasite development, such as amoeba vs circular shape rings, hemozoin formation, or differentiation of the final stages of schizogony before and after cytokinesis. Similarly, our images clearly allowed us to observed different drug-induced phenotypes such as a food vacuole defect, arrest in mid schizogony, block of egress or the pyknotic phenotype due to parasite death. Finally, we anticipate that other commercially available dyes that stain specific organelles in the cells, such as lysotracker or mitotracker, could be combined with the 132A RNA dye to obtain more detailed and specific phenotypes. Importantly, our automatic image analysis workflow can directly be applied to any samples where the nucleus and cytosol have been stained, either using different fluorescent dyes than the ones used in this study, or using fluorescent protein makers.

## Materials and Methods

### Parasite cultures and compound treatment

3D7, 1G5 and DPAP3cKO *P. falciparum* parasite lines were cultured in RPMI 1640-based media supplemented with Albumax (Invitrogen) as previously described^41^. Anonymized human blood to culture malaria parasites was purchased from the United Kingdom National Health System Blood and Transplant Special Health Authority. No ethical approval is required for its use. To obtain tightly synchronized cultures, schizonts were purified via Percoll gradient and incubated under shaking conditions with RBCs for 1h. Unruptured schizonts were removed by Percoll gradient and ring stage cultures obtained after sorbitol treatment. Parasitemia was then adjusted to 3% and cultures treated with DMSO or 100 nM RAP for 3h, washed with RPMI, and cultured in 5 identical 96-round bottom well plates (200uL of culture at 1% hematocrit) for 76h. Each plate contained 3 replicates of each culture. Twenty microliter aliquots were collected from each well at different time points, fixed with 20 μL of 8% PFA and 0.04% glutaraldehyde in PBS for 1h, followed by the addition of 200 μL of PBS and storage of samples at 4°C (0.1% hematocrit). To avoid parasite cultures from slowing down due to the collection of samples, we rotated from which plate each time point sample was collected.

Compound treatment was performed in tightly synchronized cultures of 3D7 parasites. Treatment with CQ (100nM), E64 (10 μM), and NMTI (200 μM) was performed at 4 hpi and samples collected at 48 hpi. For C2 treatment (1 μM), cultures were treated at 42 hpi and sample collected at 65 hpi. Parasites were cultured in 96-well plates with nine technical replicates for each treatment.

The A1-H.1 *P. knowlesi* strain was cultured in RPMI-based media supplemented with horse serum as previously described^31^. To obtain a tightly synchronized culture of *P. knowlesi* parasites, schizonts were first purified by Percoll gradient and cultured under shaking conditions for 4h in the presence of RBCs, which allowed the most matured schizonts to rupture and merozoites to invade RBCs. A second Percoll gradient was then used to purify the remaining schizonts and these were incubated with RBC in the presence of 1 μM C2 for 4h. The culture was then washed with RPMI and incubated under shaking condition for 2h in media. A final Percoll gradient was performed to remove as many unruptured schizonts as possible, and the resulting ring-stage parasite culture plated in 96-well plates. Samples were collected at different time points in triplicate as described above for *P. falciparum*.

### Staining protocol

For flow cytometry analysis, 10 μL of 10 μM 132A and 5 μg/mL Hoechst was added to 40 μL of sample and incubated at 37°C for 30 min in 384-well plates. For microscopy analysis, 50 μL of sample was treated with 50 μL of 4 μM 132A, 2 μg/mL Hoechst, and 0.2 μg/mL of WGA-647, transferred to a glass-bottom 96-well plate and incubated at 37°C for 30 min, which is sufficient time to allow RBCs to settle at the bottom of the well.

### Flow Cytometry Analysis

Samples stained as described above were run on a Fortessa flow cytometer (BD Bioscience) equipped with a plate autosampler using the high throughput mode. Samples were mixed three times, and 10 uL of sample were injected into the flow cytometer at 0.5 uL/s (5000-10000 RBCs per samples). FlowJo was used to analyze the flow cytometry data. RBCs were gated according to their size using the forward and side light scattering channels. iRBCs were distinguished from uRBCs as the population positive for DNA staining. For each time point, the median DNA and RNA signal intensities of iRBCs were divided by those of the uRBCs population. This normalization allowed us to obtain consistent values across different flow cytometers. Finally, these values were normalized according to the initial values obtained at 1 hpi. Note that during egress and invasion, mature schizonts and newly iRBCs were gated as two distinct populations based on the difference in DNA and RNA staining.

### Fluorescence Microscropy

Images were collected on an Olympus IX83 inverted microscope using a PLAPON O 60/1.42 objective lens and a Hamamatsu ORCA-Flash 4.0 sCMOS camera (pixel size 6.5um). Acquisition software was Olympus CellSens dimention v1.6 with the well plate navigator. A drop of immersion oil was manually dispensed under each well and twenty-five images per well were collected in four channels; BF (differential interference contrast), Hoechst (λ_ex_ = 390/18 nm, λ_em_ = 440/40nm), RNA (λ_ex_ = 560/25nm, λ_em_ = 607/36nm) and WGA-647 channels (λ_ex_ = 650/20 nm, λ_em_ = 692/40 nm). This resulted in the imaging of about 200 iRBCs per well.

### Automated analysis of fluorescence microscopy images

Images acquired from the Olympus IX83 microscope were converted to separated tif images using a Fiji^42^ script. Separate tif files were used as set of images to run a CellProfiler^43^ pipeline that identifies every single iRBC in the imaged field as objects containing DNA (Hoechst) and RNA (132A) signal within the confines of the RBC membrane signal (WGA-647). CellProfiler^44^ then measured intensity, size, shape, granularity, and texture from the raw images under the selected object in the four imaged channels (DIC, RNA, DNA, WGA-647) and exports these measurements as a CSV file. In addition, cell profiler exports thumbnail images of each of the identified objects (iRBCs) as an SQLite Database that can be use with CellProfiler Analyst^44^. These thumbnail images show either a merged image of the fluorescent channels (DNA, RNA, and WGA-647) or a merged image of the DIC, RNA and DNA channels.

An R script was then created to import the CSV file generated by CellProfiler to perform the cluster analysis. This script imports and organize the data from the CellProfiler pipeline, allows visualization of the data, performs the t-SNE multiple component analysis, and graph the results of this cluster analysis. The script and its output can be seen in the Final002.nb.KNOWLESI.html file in the supplementary materials. The supplementary file “Complete pipeline with summary notes.docx” provides a step-by-step summary of this automated analysis of the images.

## Acknowledgements

This work was supported by a Welcome Trust and Royal Society Sir Henry Dale Fellowship 099950. This work was also supported by the Francis Crick Institute which receives its core funding from Cancer Research UK (FC001043), the UK Medical Research Council (FC001043), and the Wellcome Trust (FC001043). For the purpose of Open Access, the author has applied a CC BY public copyright licence to any Author Accepted Manuscript version arising from this submission.

