## Supplementary Methods for "High Content and High Throughout Phenotypic Assay for the Hourly Resolution of the Malaria Parasite Erythrocytic Cycle"

### Imaging:

Cells were imaged on an Olympus IX83 microscope with a x60/1.42 oil immersion objective onto a Flash 4.0 sCMOS. CellSens dimension software using well plate navigator module to automate the acquisition.

### Preprocessing:

Acquired Olympus .vsi files were batch converted into separate .tif images (for the benefit of CellProfiler Anylist cell display) using a Fiji script.
"IJMacro_BatchVSI_ForCP_4C_x60_00.ijm".

### CellProfiler:

Separate .tifs files were then used as sets of images and run through a CellProfiler Pipeline "DBx60 complete pipeline with notes.cpproj". Data was saved from CellProfiler as sets of .csv files to be processed in R. In addition an SQLite database was created to be used for preliminary analysis in CellProfiler Analyst.

CellProfiler pipeline has 24 active steps in it and the pipeline can be subdivided into four main tasks. Standard CellProfiler modules were used and settings established empirically.

#### Load files

- 1-4 Load the relevant files, extract metadata from their names and group them.

#### Find infected red blood cells (iRBCs)

- 5 Find RBCs [Objects - MembsSmooth]
- 6 Mask DNA images with MembsSmooth which are the found RBCs [Images - MaskDNA]
- 7 Enhance DNA signal in MaskDNA with speckle detector [Images – EnhancedNuclei]
- 8 Find nuclei in EnhancedNuclei images [Objects – Nuclei]
- 9 Filter Nuclei objects to exclude noisy cells where far too many nuclei were found (<100) [Objects - NucleiFiltered]
- 10 Merge touching Nuclei in NucleiFiltered (probably the same nucleus anyway) [Objects – MergedNuclei]
- 12 Relate MembsSmooth (parent objects) to MergedNuclei (child objects) to make new object set [Objects – InfectedMembs]
- 13 Filter InfectedMasks to exclude those with greater than 20 nuclei [Objects – InfectedMasks]

#### Find RNA and relate to iRBCs

- 16 In RNAdye images find RNA objects [Objects – RNAObjects]
- 17 Relate InfectedMasks (parent) to RNAObjects (child) [WelatedRNAObjects]
- 18 Merge touching RNAObjects [Objects – MergedRNAObjects]
- 19 Relate InfectedMasks to MergedRNAObjects [Objects - WelatedRNAObjects]

#### Make measurements

- 22-27 Make Measurements of intensity, size and shape, granularity and texture from raw images under selected objects from above.

#### Save measurements and database

- 34 Save and export measurements as .csv
- 35 Save and export SQLite Database and properties file for use with CellProfiler Analyst.

### Analysis in R

An R script was created to import and process data from the CellProfiler .csv files. It can be broadly divided into three sections.

#### Import and organise data from CellProfiler pipeline:

Measurements from the InfectedMasks objects (iRBC) are brought together with size and shape measurements of related RNAObjects. In addition, extra metadata from well.defs.csv was added to relate other experimental variables to the data.

#### Sanity checks on assembled data:

Outlying data can be identified and excluded here. These images tend to be ones where there are bits of dust or detritus that contributes to very high signal, in one or more channels, to nearby RBCs. Once troublesome wells or images are identified then the raw data can be examined and decisions made to exclude them. As part of these sanity checks classifications can be brought in from CellProfiler Analyst. This is highlighted with a test classification represented in the Class column.

#### t-SNE and cluster analysis:

From the organised data set created above a refined version is made. This may just have Knowlesi data (or other as stipulated by variable cond1.choice). Also, columns of data are chosen; here most are included.

Rtsne is performed using the variables defining perplexity, learning and max Iterations which were determined empirically based on achieving good stability of the error [40,500 and 5000 respectively].

Clustering was then performed on the reduced dimension data. Analysis to determine the recommended number of clusters is included but we tried 12 clusters to match the distinct growth phases of the parasites.

#### Graphing:

After 3D t-SNE and clustering data was plotted in various ways showing distribution of the clusters with respect to Well, DNA and RNA intensity and by their relative distribution over time.

### Data export:

Files created and exported include those needed to retrieve t-SNE clustered images from the SQLite database created by CellProfiler and visualised and or exported from CellProfiler Analyst classifier. A static copy of clustered t-SNA data was also saved to be reused for the sake of consistency of graphing as there is an inherent variability from one run to the next.
