## Supplementary Methods for "High Content and High Throughout Phenotypic Assay for the Hourly Resolution of the Malaria Parasite Erythrocytic Cycle"


Code 

- Show All Code
- Hide All Code
- Download Rmd

### High Content and High Throughout Phenotypic Assay for the Hourly Resolution of the Malaria Parasite Erythrocytic Cycle


#### R processing script for RBC Data coming from CellProfiler.

This code collates exported .csv files from the CellProfiler pipeline DBx60 complete pipeline finessed002.ccproj and classifications from CellProfiler Analyst. Used output is intensity, shape and texture measurements from the identified infected red blood cells (iRBC).

Briefly: Cells were imaged on an Olympus IX83 microscope with a x60/1.42 oil immersion objective onto a Flash 4.0 sCMOS. CellSens dimension software using well plate navigator module to automate the acquisition (see materials and methods). Acquired Olympus .vsi files were batch converted into separate .tif images (for the benefit of CellProfiler Anylist cell display) using a Fiji[1] script. “IJMacro\_ED\_BatchVSI\_ForCP\_4C\_x60\_00.ijm”. Separate .tifs were then used as sets of images and run through a CellProfiler Pipeline “DBx60 complete pipeline finessed002with notes.cpproj”. Data was saved from CellProfiler as sets of .csv files to be used in the following R script and in addition an SQLite database to be used for preliminary analysis in CellProfiler Analyst[3].

Appendex at the end describes a module by module summary of the CellProfiler Pipeline.

[1]: Schindelin, J.; Arganda-Carreras, I. & Frise, E. et al. (2012), “Fiji: an open-source platform for biological-image analysis”, Nature methods 9(7): 676-682, PMID 22743772, doi:10.1038/nmeth.2019 (on Google Scholar)  
[2]: Kamentsky L, Jones TR, Fraser A, Bray M, Logan D, Madden K, Ljosa V, Rueden C, Harris GB, Eliceiri K, Carpenter AE (2011). Improved structure, function, and compatibility for CellProfiler: modular high-throughput image analysis software. Bioinformatics 2011/ doi. PMID: 21349861 PMCID: PMC3072555]  
[3]: Jones TR, Carpenter AE, Lamprecht MR, Moffat J, Silver S, Grenier J, Root D, Golland P, Sabatini DM (2009) Scoring diverse cellular morphologies in image-based screens with iterative feedback and machine learning. PNAS 106(6):1826-1831/doi: 10.1073/pnas.0808843106. PMID: 19188593 PMCID: PMC2634799

#### Import and organise data from CellProfiler pipeline

Get well and plate information from Plate master file, Well\_Defs.csv.


```
well.defs <- read.csv("Well_Defs.csv", header = T)
#Set working directory variable
WorkDir <- "/data"
```


Here are the source raw files from Cell Profiler being used below. First is data from the infected red blood cells (infected on the basis of DNA objects found) Upon importing the numbers represent the number of rows and columns. FIles used for analysis are listed. Figures indicate the size of the datafram in rows and columns.


```
##**************************************************************##
## Makes a data table from the output files from Cell Profiler  ##
##**************************************************************##
read_table_filename <- function(filename){
  ret <- read.csv(filename,header = T)
  ret$Source <- filename #EDIT
  ret
}
filenames <- list.files(pattern="InfectedMasks.csv", full.names = TRUE,recursive = TRUE)
filenames
```


```
[1] "./data/EDPlate/MyExpt_InfectedMasks.csv"             "./data/Time Course Plate 1/MyExpt_InfectedMasks.csv"
```


```
InfectedCells <- ldply(filenames, read_table_filename)

InfectedCells$Source <- gsub(".csv","", InfectedCells$Source)
InfectedCells$Source <- gsub("./","", InfectedCells$Source)
InfectedCells$Source <- as.factor(InfectedCells$Source)
dim(InfectedCells)
```


```
[1] 7686  408
```


Now comes measurements from merged RNA detections as described in the CellProfiler pipeline.  
Data frame dimensions as row and column.


```
##**********************************************************##
## Aims to extract area measurements from the RNA Objects   ##
##**********************************************************##
filenames <- list.files(pattern="MergedRNAObjects.csv", full.names = TRUE,recursive = TRUE)
filenames
```


```
[1] "./data/EDPlate/MyExpt_MergedRNAObjects.csv"             "./data/Time Course Plate 1/MyExpt_MergedRNAObjects.csv"
```


```
MergedRNA <- ldply(filenames, read_table_filename)

MergedRNA$Source <- gsub(".csv","", MergedRNA$Source)
MergedRNA$Source <- gsub("./","", MergedRNA$Source)
MergedRNA$Source <- as.factor(MergedRNA$Source)

dim(MergedRNA)
```


```
[1] 36122    82
```


Here we summarize some of the RNA data per Infected Cell so we can then add it to the iRBC data above


```
RNAtab<-ddply(ddply(MergedRNA, c('Source',
                                 'Metadata_Plate',
                                 'Metadata_Vers',
                                 'Metadata_Well',
                                 'ImageNumber',
                                 'Parent_InfectedMasks'),
                    function(x) c( numRNA=length(x$AreaShape_Area),
                                   MeanAreaRNA=mean(x$AreaShape_Area),
                                   TotAreaRNA=sum(x$AreaShape_Area) )),
              .(Metadata_Well))
colnames(RNAtab)[colnames(RNAtab)=="Parent_InfectedMasks"] <- "ObjectNumber"
#dim(RNAtab)
```


Now we take the summary table of RNA objects and in it with the Infected Cells data from above. Infected cells where no RNA objects were detected have their summary values set to 0 (rather then ).In addition some of the factors used later on (Well number and timepoint) are ordered more sensibly. Again after this chunk numbers are a check showing rows and columns in the data frame.


```
mergeInfectedCells <- join(InfectedCells, RNAtab[,c(5:9)], by= c('ImageNumber','ObjectNumber'))
mergeInfectedCells$MeanAreaRNA[is.na(mergeInfectedCells$MeanAreaRNA)] <- 0
mergeInfectedCells$TotAreaRNA[is.na(mergeInfectedCells$TotAreaRNA)] <- 0
mergeInfectedCells$numRNA[is.na(mergeInfectedCells$numRNA)] <- 0
#names(mergeInfectedCells)[duplicated(names(mergeInfectedCells))]# to find duplicated column names
mergeInfectedCells <- join(mergeInfectedCells, well.defs, by = c('Metadata_Plate', 'Metadata_Well'))
#Makes sure these variables apperar in a sensible order.
mergeInfectedCells$Metadata_Well <- factor(mergeInfectedCells$Metadata_Well, levels =  c ("B2","B3","B4","B5","B6","B7","B8","B9","B10","B11","C2","C3","C4","C5","C6","C7","C8","C9","C10","C11","D2","D3","D4","D5","D6","D7","D8","D9","D10","D11","E2","E3","E4","E5","E6","E7","E8","E9","E10","E11","F2","F3","F4","F5","F6","F7","F8","F9","F10","F11"))
mergeInfectedCells$time <- factor(mergeInfectedCells$time, levels = c("0","1","3","4","5","7","8","9","11","12","13","14","15","16","17","18","19","20","21","22","23","24","25","26","27","28","29","30","31","32","33","34","35","36","37","38","39","40","41","42","44","46","48","50","52","54","56","58","60","68","78"))
mergeInfectedCells$Unique.im.id <- with(mergeInfectedCells,  paste(Metadata_Plate, Metadata_Well,  Metadata_Image, Metadata_Vers, sep = "_"))
dim(mergeInfectedCells)
```


```
[1] 7686  416
```

#### Sanity checks on assembled data.

Initally simply plotting the mean DNA intensity per object, per image, per plate. Outlying spikes should appear obvious. Dot plot to show outlying DNA intensity per image. This is often due to rubbish in these images. I looked at these outliers by eye and indeed they are due to anomalies within the images like dust, bubbles etc.  
Note the change in y scale.


```
 #Dot plot to show outlying DNA intensity per image. This is often due to rubbish in these images
p1 <- ggplot(mergeInfectedCells, aes(ImageNumber, Intensity_MeanIntensity_DNA, colour = Metadata_Well)) +
  geom_point(size = .5) +
  theme(legend.position = "FALSE", axis.text.x = element_text(angle = 45, vjust = 0.5),text = element_text(size=10))+
  facet_wrap(.~Metadata_Plate, scales = 'free_x')
#List of excluded images, based no junk within the image. 
#Format is Platename, Well, Image, Version, separated by _.
exclude.DNA <- c("EDPlate_C7_34_1", "EDPlate_C7_35_1","EDPlate_C7_42_1","EDPlate_C7_49_1", "Time Course Plate 1_C4_13_1","Time Course Plate 1_G8_11_1", "Time Course Plate 2_C7_6_1", "Time Course Plate 2_D11_11_1", "Time Course Plate 2_E10_14_1", "Time Course Plate 2_E10_15_1", "Time Course Plate 2_F2_8_1")
exclude.RNA <- c("EDPlate_C7_27_1")
exclude.uq.im.id <- append(exclude.DNA, exclude.RNA)

`%nin%` <- Negate(`%in%`) #Quick function to help following exclusion
mergeInfectedCells <-  mergeInfectedCells[ mergeInfectedCells$Unique.im.id %nin% exclude.uq.im.id, ]

p2 <- ggplot(mergeInfectedCells, aes(ImageNumber, Intensity_MeanIntensity_DNA, colour = Metadata_Well)) +
  geom_point(size = .5) +
  theme(legend.position = "FALSE", axis.text.x = element_text(angle = 45, vjust = 0.5),text = element_text(size=10))+
  facet_wrap(.~Metadata_Plate, scales = 'free_x')

grid.arrange(p1,p2,nrow = 1, top=textGrob("Sanity check plots before and after exclusion - by image number", gp=gpar(fontsize=14)))
```


Data is trimmed further to only include the timelapse data for P.Falciparum or P.Knowlesi. Plots are of the respective strains over time (by well number).


```
#Make a duplicate copy for use later if necessary
mergeInfectedCells2 <- mergeInfectedCells
#Choose only those rows that are described as Knowlesi or Falciparum in the Well_Defs.csv
mergeInfectedCells <- mergeInfectedCells[mergeInfectedCells$Cond1 == "Falciparum" | mergeInfectedCells$Cond1 == "Knowlesi",]

p1 <-ggplot(mergeInfectedCells[!is.na(mergeInfectedCells$Cond1),], aes(Metadata_Well, Intensity_MedianIntensity_RNAdye, colour = Metadata_Well)) +
    geom_jitter(size = 0.2) +
    theme(legend.position = "FALSE", axis.text.x = element_text(angle = 90, vjust = 0.5),text = element_text(size=8))+
    facet_grid(Cond1~., scales = 'free')

p2 <- ggplot(mergeInfectedCells[!is.na(mergeInfectedCells$Cond1),], aes(Metadata_Well, Intensity_MedianIntensity_DNA, colour = Metadata_Well)) +
    geom_jitter(size = 0.2) +
    theme(legend.position = "FALSE", axis.text.x = element_text(angle = 90, vjust = 0.5),text = element_text(size=8))+
    facet_grid(Cond1~., scales = 'free')
grid.arrange(p1,p2,nrow = 1, top=textGrob("Sanity check plots timelapse data only, RNA and DNA intensity", gp=gpar(fontsize=14)))
```


```
mergeInfectedCells <- mergeInfectedCells2
```


Using manual classifications from CellProfiler Analyst TrainingSet file. This are what cell profiler analyst uses to predict classifications for the rest of the dataset (see below). We dont necessarily use these classifications but its an easy way to identify these first training examples on subsequent plots.


Trained classifications from CellProfiler Analyst CLASS\_PerObj files. Based on manual training set (above). Brings these definitions into our master data set “mergeInfectedCells”. With these variables we can see how the CellProfiler Analyst classifications line up with clusters identified in subsequent analysis. This list is generated from the CPA/classifier module.


```
filenames <- list.files(pattern="my_table.csv", full.names = TRUE,recursive = TRUE)
#filenames 
CLASSes <- ldply(filenames, read_table_filename)
colnames(CLASSes)[colnames(CLASSes)=="InfectedMasks_Number_Object_Number"] <- "ObjectNumber"
colnames(CLASSes)[colnames(CLASSes)=="class"] <- "Class"
CLASSes$Class<-as.factor(CLASSes$Class)
##********************************************************************************************##
##Use the next line if no class table is available
#CLASSes <- data.frame(ImageNumber = mergeInfectedCells$ImageNumber, ObjectNumber = mergeInfectedCells$ObjectNumber, Source = mergeInfectedCells$Source, Class = "NoneAssigned" )
##********************************************************************************************##
dim(CLASSes)
```


```
[1] 39932     5
```


```
mergeInfectedCells <- join(mergeInfectedCells, CLASSes, type = "left", by= c('ImageNumber','ObjectNumber'))
mergeInfectedCells <- mergeInfectedCells[,!duplicated(names(mergeInfectedCells))]
mergeInfectedCells$Class<-as.factor(mergeInfectedCells$Class)
summary(mergeInfectedCells$Class)
```


```
  BigRNA negative Schisont SmallRNA 
    2551      340      766     3932
```


```
dim(mergeInfectedCells)
```


```
[1] 7589  419
```


This next bit gets rid of any duplicates but also uses a modified list of objects from Edgar (EDmod\_list.csv). In essence this is a refinement of the trained set, checked by eye and unusual objects removed eg multiple infections.


```
mergeInfectedCells <- mergeInfectedCells[,!duplicated(names(mergeInfectedCells))]
dim(mergeInfectedCells)
```


```
[1] 7589  419
```


```
##*********************************************************************************************##
## Using Modified list, manually sorted from output of cluster analysis (from ED). 
##*********************************************************************************************##
filenames <- list.files(pattern="EDmod_list.csv", full.names = TRUE,recursive = TRUE)
read_table_filename <- function(filename){
  ret <- read.csv(filename,header = T, sep=",")
  #ret$Source <- filename #EDIT
  ret
}

EDmod <- ldply(filenames, read_table_filename)
mergeInfectedCells <- join(mergeInfectedCells, EDmod, by= c('ImageNumber','ObjectNumber'))
mergeInfectedCells$EDmod_Cluster[is.na(mergeInfectedCells$EDmod_Cluster)] <- "XX"
dim(mergeInfectedCells)
```


```
[1] 7589  420
```


```
mergeInfectedCellsSmall <- mergeInfectedCells[,c(1:14,35:98,126:407,409:414)]
dim(mergeInfectedCellsSmall)
```


```
[1] 7589  366
```


This just saves a copy of the data calculated so far into the out directory. It allows us to jump into the script halfway though.


```
##*********************************************************************************************##
## Save....Objects
##*********************************************************************************************##

write.csv(mergeInfectedCells,"out/mergeInfectedCells.csv")
write.csv(mergeInfectedCellsSmall,"out/mergeInfectedCellsSmall.csv")
```

#### t-SNE tests.

This is the beginning of the R t-SNE test for the above assembled data. Can come in at this point if mergeInfectedCells.csv is loaded


```
##********************************************************************************************##
##Shortcut to mergeInfectedCells file
##********************************************************************************************##
#mergeInfectedCells <- read.csv("out/mergeInfectedCells.csv")
#mergeInfectedCells <- mergeInfectedCells[,-1]
#This line removes ED manually selected troublesome images
#mergeInfectedCells <- mergeInfectedCells[mergeInfectedCells$EDmod_Cluster != "XX",]
mergeInfectedCells2 <- mergeInfectedCells
```


Here we create an object for the cluster analysis (called all.tsne). This can be further filtered to only include one or other of the plasmodium strains.


```
##********************************************************************************************##
##Create an object for the cluster analysis (called all.tsne)
##********************************************************************************************##
#all.tsne <- mergeInfectedCells[,c(-3,-6)] # This would be everything from all plates
#variable to choose data from one paracyte or the other 
cond1.choice <- "Knowlesi"
#cond1.choice <- "Falciparum"

all.tsne <- mergeInfectedCells[mergeInfectedCells$Cond1 == cond1.choice,]
#removes columns that have NA's
all.tsne <- all.tsne[,c(-3,-6)] 
#removes weird rows
all.tsne <- all.tsne[!is.na(all.tsne$ImageNumber),]
dim(all.tsne)
```


```
[1] 3060  418
```


```
#ddply(ddply(all.tsne, c('Metadata_Plate','Metadata_Well','Class','Cond1'), function(x) c(number.Class=length(x$Class))),.(Metadata_Plate,Metadata_Well))
```


Sanity check table showing a summary of the Class types per well


```
#Summary table for selected samples showing the number of each class per well
head(ddply(ddply(all.tsne, c('Metadata_Plate','Metadata_Well','Class','Cond1'), function(x) c(number.Class=length(x$Class))),.(Metadata_Plate,Metadata_Well)))
```


Make an object that just has measurements in it for t-SNE analysis (ED.data) and process this with using 2 and 3 dimensions. Choice of perplexity, learning and maximum Iterations was determined empirically by adjusting the respective setings but a perplexity of 40, learning of 500 and maximum Iterations of 5000 seemed to give a good compromise between processing time and clustering. The graph of 2D t-SNE analysis shows colours indicating the EDmod\_clusters (clusters from a previous analysis of the same data). These were then manually reviewed and images selected showing how the excluded/ambiguous images are more prevalent in some areas than others (XX=White), possibly due to having measurements that are otherwise quite similar to certain stages of the malaria life cycle. The following analysis was, however, carried out on the entire dataset as it would be for any new samples.


Clustering performed on the tsne3D output. It is possible to change the number of clusters here. We chose a number to try and match the number of malaria stages represented in the timelapse experiment.


```
#Set number of clusters to be used
clusts <- 12

#3D clustering
x <- tsne3D[,c(3:5)]
#head(x)

library(factoextra)
library(NbClust)

fviz_nbclust(x, kmeans)
```


```
#fviz_nbclust(x, kmeans, method = "wss")
#fviz_nbclust(x, kmeans, method = "silhouette")
#res <- NbClust(data = x,  distance = "euclidean", min.nc = 2, max.nc = 15, method = "complete")
#res$Best.nc


hc = hclust(dist(x), method = "ward.D")
hc
```


```
Call:
hclust(d = dist(x), method = "ward.D")

Cluster method   : ward.D 
Distance         : euclidean 
Number of objects: 3060
```


```
cluster_grps_12<- cutree(hc, k = clusts) 

k <- kmeans(x, clusts, nstart=25, iter.max=1000)
#k
new = cbind(x,k.cluster = k$cluster)
new = cbind(new,hc = cluster_grps_12)
#tail(new)
#head(new)
#plot3d(new, col=new$k.cluster)

tsne3D$k.cluster3D <- new$k.cluster
tsne3D$k.cluster3D <- as.factor(tsne3D$k.cluster3D)
tsne3D$hc.cluster3D <- new$hc
tsne3D$hc.cluster3D <- as.factor(tsne3D$hc.cluster3D)

#Output the current version of the t-SNE performed in 3D.
write.csv(tsne3D, paste("out/", cond1.choice,"_tsne3D.csv", sep = ""))
```


Here is a # out entry point for a specified t-SNE and clustering run. The nature of the analysis is that there is variability each time it is run so this allows entry at a static point for graphing.


Starting point for graphing. Colours an levels (order) of clusters can be specified and these were informed here at the end of the analysis by staging malarias in iRBC’s in the respective clusters (for consistancy of graphing).


```
#For the benefit of getting the colours in the right order when using scatter3D it must be in the order of the levels.
EDcols2.k <- c( "1" = "#12100B", "2" = "#E52613", "3" = "#FFD877", "4" = "#414F9D", "5" = "#961914", "6" =  "#B7B7B7", "7" = "#61B22F", "8" = "#008938","9" = "#C75D9F", "10" = "#F39000" , "11" =  "#2C2B7B", "12" =  "#90D4F6")
EDcols.K <- c( "A1" = "#12100B", "B2" = "#E52613", "C3" = "#FFD877", "D4" = "#414F9D", "E5" = "#961914", "F6" =  "#B7B7B7", "G7" = "#61B22F", "H8" = "#008938","I9" = "#C75D9F", "J10" = "#F39000" , "J11" =  "#2C2B7B", "L12" =  "#90D4F6")

EDcols2.hc <- c( "1" = "#12100B", "2" = "#961914", "3" = "#414F9D", "4" = "#C75D9F", "5" = "#E52613", "6" =  "#2C2B7B", "7" = "#61B22F", "8" = "#F39000","9" = "#FFD877", "10" = "#008938" , "11" =  "#90D4F6", "12" =  "#B7B7B7")
EDcols.HC <- c( "A1" = "#12100B", "B2" = "#961914", "C3" = "#414F9D", "D4" = "#C75D9F", "E5" = "#E52613", "F6" =  "#2C2B7B", "G7" = "#61B22F", "H8" = "#F39000","I9" = "#FFD877", "J10" = "#008938" , "K11" =  "#90D4F6", "L12" =  "#B7B7B7")

ggplot(tsne3D) + geom_point(aes(x=y, y=z, color=factor(hc.cluster3D)), size = 1) + ggtitle(paste(cond1.choice,": tsne3D kmeans", clusts))+ scale_colour_manual(values = EDcols2.hc) #+facet_grid(k.cluster3D~Metadata_Well)

require(rgl)
require(car)
#palette(crick_pal()(clusts))
palette(EDcols2.hc)
```


```
#palette(rainbow(clusts)) # Or use your own palette...

scatter3d(x = tsne3D$x, y = tsne3D$y, z = tsne3D$z, groups = factor(tsne3D$hc.cluster3D),surface.col = 1:clusts,
          surface=FALSE, ellipsoid = FALSE, ellipsoid.alpha = 0.01, classLabel = c("sommething","something"))
rgl.snapshot(filename = paste("out/",cond1.choice, "_nohull_tsne3D", fmt = ".png", sep=""))
clear3d()
scatter3d(x = tsne3D$x, y = tsne3D$y, z = tsne3D$z, groups = factor(tsne3D$hc.cluster3D),surface.col = 1:clusts,
          surface=FALSE, ellipsoid = TRUE, ellipsoid.alpha = 0.08, bg.col=c( "white"),)

rgl.snapshot(filename = paste("out/",cond1.choice, "_NEW_hull_tsne3D", fmt = ".png", sep = ""))
```


3D scatterplots are produced with this code and follows is a snapshot of one of those. It is interactive when code is run.

First steps in segmentation

Plots of wells versus clusters so we can start to see change over time. Firstly we see these as a 2D plot (from the 3D data). Some wells/timepoints were excluded due to imaging anomalies.  
By well scatterplots showing progression of different clusters throughout the experiment


```
tsne3D002 <- tsne3D[tsne3D$Metadata_Well != "D2",]
tsne3D002 <- tsne3D002[tsne3D002$Metadata_Well != "D3",]
tsne3D002 <- tsne3D002[tsne3D002$Metadata_Well != "D8",]

tsne3D002$EDmod_cluster[is.na(tsne3D002$EDmod_cluster)] <- "Junk"

#p1 <- ggplot(tsne3D002) + geom_point(aes(x=y, y=z, color=Class), size = 0.5) +facet_wrap(.~Metadata_Well)+ ggtitle(paste(cond1.choice,": tsne3D CPA classes")) + theme(legend.position = "Null") + theme(plot.title = element_text(size = 11, face = "bold"))
p2 <- ggplot(tsne3D002) + geom_point(aes(x=y, y=z, color=factor(hc.cluster3D)), size = 0.5) +facet_wrap(.~Metadata_Well)+ ggtitle(paste(cond1.choice, ": tsne3D hc ", "Clusters: ", clusts)) + theme(legend.position = "Null")+ scale_colour_manual(values = EDcols2.hc) + theme(plot.title = element_text(size = 11, face = "bold"))
p2
```


```
#grid.arrange(p1, p2, nrow = 1)
```


Code to just arrange data into an object for following plots. Troublesome wells excluded.


```
v = factor(tsne3D$k.cluster3D)
levels(v) = c("A1","B2","C3","D4","E5","F6","G7","H8","I9","J10","K11","L12")
vv = factor(tsne3D$hc.cluster3D)
levels(vv) = c("A1","B2","C3","D4","E5","F6","G7","H8","I9","J10","K11","L12")
Rtsne3DTrained <- data.frame(ImageNumber=all.tsne$ImageNumber, ObjectNumber=all.tsne$ObjectNumber,  Metadata_Well = all.tsne$Metadata_Well, tsne1 = tsne3D$x, tsne2 = tsne3D$y , tsne3 =tsne3D$z , k.cluster3D =  v, hc.cluster3D =  vv)
Rtsne3DTrained <- join(all.tsne, Rtsne3DTrained, by = c("ImageNumber", "ObjectNumber","Metadata_Well"))
#dim(Rtsne3DTrained)

Rtsne3DTrained002 <- Rtsne3DTrained[Rtsne3DTrained$Metadata_Well != "D2",]
Rtsne3DTrained002 <- Rtsne3DTrained002[Rtsne3DTrained002$Metadata_Well != "D3",]
Rtsne3DTrained002 <- Rtsne3DTrained002[Rtsne3DTrained002$Metadata_Well != "D8",]
```


Scatter plots coloured and faceted by hc.cluster. Median intensities are shown for DNA and RNAdye.


```
orderK <- c("G7","H8","L12","D4","K11","C3","J10","B2","E5","I9","A1","F6")
orderHC <- c("G7","J10","K11","C3","F6","I9","H8","E5","B2","D4","A1","L12")
orderEDmod <- c("B2","I9","C3","E5","L12","J10","K11","D4","G7","A1","F6","H8")
Rtsne3DTrained002 <- arrange(transform(Rtsne3DTrained002,
             hc.cluster3D=factor(hc.cluster3D,levels=orderHC)),hc.cluster3D)
Rtsne3DTrained002 <- arrange(transform(Rtsne3DTrained002,
             k.cluster3D=factor(k.cluster3D,levels=orderK)),k.cluster3D)
Rtsne3DTrained002 <- arrange(transform(Rtsne3DTrained002,
             EDmod_Cluster=factor(EDmod_Cluster,levels=orderEDmod)),EDmod_Cluster)

#p7 <- ggplot(Rtsne3DTrained002, aes(x = Intensity_MedianIntensity_DNA, y = Intensity_MedianIntensity_RNAdye, colour = Class)) + geom_point(alpha = 0.6, size = 0.5) + theme_bw() + ggtitle(paste(Rtsne3DTrained002$Cond1,"Facet by EDmod_clusters", "Clusters: ", clusts)) + facet_wrap(.~EDmod_Cluster)#+ theme(legend.position = "false")
#p7
p8 <- ggplot(Rtsne3DTrained002, aes(x = Intensity_MedianIntensity_DNA, y = Intensity_MedianIntensity_RNAdye, colour = hc.cluster3D)) + 
  geom_point(alpha = 0.6, size = 0.5) + theme_bw() + #coord_cartesian(xlim = c(0,0.065), ylim = c(0,0.065)) +
  ggtitle(paste(Rtsne3DTrained002$Cond1,"Facet by EDmod_clusters", "HC Clusters: ", clusts)) + facet_wrap(.~hc.cluster3D)+ scale_colour_manual(values = EDcols.HC)+ theme(legend.position = "null")
p8
```


```
#grid.arrange( p7, p8, nrow = 1)
```


Overall median intensities of DNA and RNAdye plotted against each other for the different k or hc clusters with + and - standard error bars. Equivalent to graphs in manuscript. Note different (arbitary) cluster namesbetween k and hc types).


```
require(plotrix)
require(scales)
require(ggrepel)

z <- ddply(Rtsne3DTrained002,.(k.cluster3D),summarise,
           MedianRNA = median(Intensity_MedianIntensity_RNAdye),
           MedianDNA = median(Intensity_MedianIntensity_DNA),
           RNA.SE = std.error(Intensity_MedianIntensity_RNAdye),
           DNA.SE = std.error(Intensity_MedianIntensity_DNA))
#z
p1 <- ggplot(z,aes(x = MedianDNA,y = MedianRNA, label = k.cluster3D)) + 
  geom_point(aes(colour = k.cluster3D)) + 
  geom_text_repel(aes(label = k.cluster3D , color = k.cluster3D), size = 3.5) +
  geom_point(data = z,aes(colour = k.cluster3D)) +
  geom_errorbarh(aes(xmax = MedianDNA + DNA.SE, xmin = MedianDNA - DNA.SE, colour = k.cluster3D)) +
  geom_errorbar(aes(ymin = MedianRNA - RNA.SE, ymax = MedianRNA + RNA.SE,colour = k.cluster3D)) + 
  scale_x_continuous(trans = log2_trans(),breaks = trans_breaks("log2", function(x) 2^x),labels = trans_format("log2", math_format(2^.x)))+#(trans = 'log2' ,breaks = trans_breaks("log2", function(x) 2^x)) +
  scale_y_continuous(trans = log2_trans(),breaks = trans_breaks("log2", function(x) 2^x),labels = trans_format("log2", math_format(2^.x))) + theme(legend.position = "Null")+
  ggtitle("Colour = k.cluster3D") + scale_colour_manual(values = EDcols.K)

y <- ddply(Rtsne3DTrained002,.(hc.cluster3D),summarise,
           MedianRNA = median(Intensity_MedianIntensity_RNAdye),
           MedianDNA = median(Intensity_MedianIntensity_DNA),
           RNA.SE = std.error(Intensity_MedianIntensity_RNAdye),
           DNA.SE = std.error(Intensity_MedianIntensity_DNA))

p2 <- ggplot(y,aes(x = MedianDNA,y = MedianRNA, label = hc.cluster3D)) + 
  geom_point(aes(colour = hc.cluster3D)) + 
  geom_text_repel(aes(label = hc.cluster3D , color = hc.cluster3D), size = 3.5) +
  geom_point(data = y,aes(colour = hc.cluster3D)) +
  geom_errorbarh(aes(xmax = MedianDNA + DNA.SE, xmin = MedianDNA - DNA.SE, colour = hc.cluster3D)) +
  geom_errorbar(aes(ymin = MedianRNA - RNA.SE, ymax = MedianRNA + RNA.SE,colour = hc.cluster3D)) + 
  scale_x_continuous(trans = log2_trans(),breaks = trans_breaks("log2", function(x) 2^x),labels = trans_format("log2", math_format(2^.x)))+#(trans = 'log2' ,breaks = trans_breaks("log2", function(x) 2^x)) +
  scale_y_continuous(trans = log2_trans(),breaks = trans_breaks("log2", function(x) 2^x),labels = trans_format("log2", math_format(2^.x))) + theme(legend.position = "Null")+
  ggtitle("Colour = hc.cluster3D") + scale_colour_manual(values = EDcols.HC)

grid.arrange( p1, p2, nrow = 1)
```


```
NA
NA
```


This is about reordering the clusters newly acquired to compare with the original ones and EDmod\_Clusters. The table shows how the different clusters compare iRBC by iRBC. k and HC.clusters are actually very similar.  
First table : EDmod\_cluster (rows) versus k.cluster (cols).


```
Rtsne3DTrained002$hc.cluster3D <- factor(Rtsne3DTrained002$hc.cluster3D, levels =  c("G7","J10","K11","C3","F6","I9","H8","E5","B2","D4","A1","L12")) # order in fig6
#Sanity check
table(Rtsne3DTrained002$EDmod_Cluster,Rtsne3DTrained002$k.cluster3D)
```


```
       G7  H8 L12  D4 K11  C3 J10  B2  E5  I9  A1  F6
  B2   37  10   0   0   0   0   0   0   0   0   0   0
  I9    0 210   1   0   0   0   0   0   0   0   0   0
  C3    0   1 281   0   1   0   0   0   0   0   0   0
  E5    0   0   4 227   0   0   0   0   0   0   0   0
  L12   0   0   0   0 117   0   0   0   0   8   0   0
  J10   0   0   0  53   0 124   0   0   0   0   0   0
  K11   0   0   0   0   0  94  87   0   0   0   0   0
  D4    0   0   0   0   0   0 120 121   0   0   0   0
  G7    0   0   0   0   0   0   0 110 291   0   0   0
  A1    0   0   0   0   0   0   0   0   0 125   0   0
  F6    0   0   0   0   0   0   0   0  53   0 271   0
  H8    0   9   0   0   0   0   0   0   0   0   0  12
```


```
#Second table : EDmod_cluster (rows) versus k.cluster (cols).  
#table(Rtsne3DTrained002$EDmod_Cluster,Rtsne3DTrained002$hc.cluster3D)
#Third table is hc.clusters (rows) versus k.cluster (cols).  
#table(Rtsne3DTrained002$k.cluster3D,Rtsne3DTrained002$hc.cluster3D)
```


Density graphs of cluster prevalence over time. Either faceted separately or collectively.


```
RtsneFreqs <- Rtsne3DTrained002 %>%  
  group_by(Metadata_Well, time, hc.cluster3D) %>% 
  summarise(Freq = n())
```


```
`summarise()` regrouping output by 'Metadata_Well', 'time' (override with `.groups` argument)
```


```
RtsneFreqs2 <- RtsneFreqs %>%
  group_by(time) %>% #do calculations by siteID
  mutate(percent = Freq / sum(Freq))
#RtsneFreqs2

write.csv(RtsneFreqs2, paste("out/", cond1.choice,"HC_RtsneFreqs2.csv", sep = ""))

p1 <- ggplot(Rtsne3DTrained002, aes(time, group = hc.cluster3D, colour = hc.cluster3D, fill = hc.cluster3D)) +
  geom_density(alpha = 0.6)+
  facet_grid(hc.cluster3D~.) + 
  ggtitle(paste(cond1.choice,": tsne3D hc.cluster", clusts))+
  theme_minimal()+
  theme(legend.position = "Null")+
  theme(legend.position = "FALSE", axis.text.x = element_text(angle = 90, vjust = 0.5,size=6),text = element_text(size=10))+
  scale_colour_manual(values = EDcols.HC) +
  scale_fill_manual(values = EDcols.HC)

p2 <- ggplot(Rtsne3DTrained002, aes(time, group = hc.cluster3D, colour = hc.cluster3D, fill = hc.cluster3D)) +
  geom_density( position = "stack")+
  ggtitle(paste(cond1.choice,": tsne3D hc.cluster", clusts))+
  theme_minimal()+
  theme(legend.position = "Null")+
  theme(legend.position = "FALSE", axis.text.x = element_text(angle = 90, vjust = 0.5,size=6),text = element_text(size=10))+
  scale_colour_manual(values = EDcols.HC)+
  scale_fill_manual(values = EDcols.HC)
brks <- c(0,.25,.50,.75,1)
p3 <- ggplot(data=RtsneFreqs2,aes(x=time,y=percent,fill=hc.cluster3D)) +
  geom_bar(stat="identity") +
  scale_y_continuous(breaks = brks, labels = scales::percent(brks)) +
  scale_colour_manual(values = EDcols.HC)+
  scale_fill_manual(values = EDcols.HC)+
  theme(legend.position = "Null")+
  theme(legend.position = "FALSE", axis.text.x = element_text(angle = 90, vjust = 0.5,size=6),text = element_text(size=10))

grid.arrange( p1, p2, p3,nrow = 1)
```


Scatter plots showing DNA and RNA intensities for iRBCs over time faceted and coloured by hc.cluster.


```
p9 = ggplot(Rtsne3DTrained002, aes(time, Intensity_MedianIntensity_DNA, colour = hc.cluster3D)) + geom_jitter(size= 0.4) + facet_wrap(.~hc.cluster3D) + theme(legend.position = "blank") + ggtitle(paste(cond1.choice,"Median DNA int.", "Clusters: ", clusts)) + theme(plot.title = element_text(size = 10))+
  theme(legend.position = "FALSE", axis.text.x = element_text(angle = 90, vjust = 0.5,size=6),text = element_text(size=10))+
  scale_colour_manual(values = EDcols.HC)

p10 = ggplot(Rtsne3DTrained002, aes(time, Intensity_MedianIntensity_RNAdye, colour = hc.cluster3D)) + geom_jitter(size= 0.4) + facet_wrap(.~hc.cluster3D) + theme(legend.position = "blank") +
 ggtitle(paste(cond1.choice,"Median RNAdye int.", "Clusters: ", clusts)) + theme(plot.title = element_text(size = 10)) + theme(legend.position = "FALSE", axis.text.x = element_text(angle = 90, vjust = 0.5,size=6),text = element_text(size=10))+
  scale_colour_manual(values = EDcols.HC)

grid.arrange( p9, p10, nrow = 1)
```


```
NA
NA
```


Generation of a .csv called xxx\_RtsneTrainedSet.csv which can be imported into the classifier of CellProfiler Analyst to retrieve now clustered images. The file can be edited as the one used by the classifier to create bins is called “Class”. Once imported images from the espective classes can be exported.


```
RtsneTrainedSet2 <- data.frame(ImageNumber=Rtsne3DTrained002$ImageNumber,
                               InfectedMasks_Number_Object_Number=Rtsne3DTrained002$ObjectNumber,
                               Metadata_well = Rtsne3DTrained002$Metadata_Well,
                               Image = Rtsne3DTrained002$Metadata_Image,
                               Class = Rtsne3DTrained002$Class,
                               EDmod_cluster = Rtsne3DTrained002$EDmod_Cluster,
                               k.cluster3D = Rtsne3DTrained002$k.cluster3D,
                               hc.cluster3D = Rtsne3DTrained002$hc.cluster3D)
head(RtsneTrainedSet2)
```


```
simplified <- data.frame(Metadata_plate = Rtsne3DTrained002$Metadata_Plate,
                         Metadata_well = Rtsne3DTrained002$Metadata_Well,
                         ImageNumber=Rtsne3DTrained002$ImageNumber,
                         ObjectNumber=Rtsne3DTrained002$ObjectNumber,
                         Image = Rtsne3DTrained002$Metadata_Image,
                         Class = Rtsne3DTrained002$Class,
                         EDmod_cluster = Rtsne3DTrained002$EDmod_Cluster,
                         k.cluster3D = Rtsne3DTrained002$k.cluster3D,
                         hc.cluster3D = Rtsne3DTrained002$hc.cluster3D,
                         MeanIntensity_DNA = Rtsne3DTrained002$Intensity_MeanIntensity_DNA,
                         MeanIntensity_RNAdye = Rtsne3DTrained002$Intensity_MeanIntensity_RNAdye,
                         MedianIntensity_DNA = Rtsne3DTrained002$Intensity_MedianIntensity_DNA,
                         Mediantensity_RNAdye = Rtsne3DTrained002$Intensity_MedianIntensity_RNAdye)
head(simplified)
```


```
write.csv(RtsneTrainedSet2, paste("out/", cond1.choice,"_newRtsneTrainedSet.csv", sep = ""))
write.csv(Rtsne3DTrained002, paste("out/",cond1.choice,"_newRtsne3DTrained002.csv", sep = ""))
write.csv(simplified, paste("out/",cond1.choice,"_newSimplifiedOutput.csv", sep = ""))
```


##Appendix #CellProfiler summary steps of modules:

[ 1] [Images] Images to be loaded. Separate .tifs as use with Cell Profiler Analyst stacked images don’t display properly.

[ 2] [Metadata] Extract file variables from file name

[ 3] [NamesAndTypes] Meaningful names applied to data: BF = DIC image DNA = malaria DNA stained with Hoechst RNAdye = 132A RNA dye Membs = WGA-647 membrane marker

[ 4] [Groups] Subsets of images generated on the basis of Plate and Well

[ 5] [IdentifyPrimaryObjects] Step 1 find all red blood cells on the basis of the far red channel (WGA-647)

[ 6] [MaskImage] Mask image to membranes found

[ 7] [EnhanceOrSuppressFeatures] Enhancement of hoescht stained paracyte nuclei using a speckle detector.

[ 8] [IdentifyPrimaryObjects] FInd parcyte nuclei (that are inside membrane masks)

[ 9] [FilterObjects] Apply image filter to exclude noisy images with far too many detected nuclei (noise)

[ 10] [SplitOrMergeObjects] Merge touching nuclei (which are probably the same nuclei anyway).

[ 11] [ShrinkToObjectCenters] (disabled) From the merged nuclei generated in the previous step, reduce these to their centers. (not used)

[ 12] [RelateObjects] Relate Membranes detected to nuclei to give InfectedMembs

[ 13] [FilterObjects] Filter out membrans that have too many ‘nuclei’ associated with them. Too many is an indication of noisy images and the threshold is set quite high. Remaining membranes contain paracyte nuclei and are referred to InfectedMasks

[ 14] [RelateObjects] Relate the filtered population of membraneMasks to merged nuclei - WelatedNuclei - not used

[ 15] [MaskImage] Using the mask of infected cells as judged by DNA content, find RNAdye masks within this population to produce MaskRNA object as output.

[ 16] [IdentifyPrimaryObjects]

[ 17] [RelateObjects] Relate detected RNA objects to particular innfected masks

[ 18] [SplitOrMergeObjects] Merge touching RNA objects

[ 19] [RelateObjects] Relate merged RNA objects to InfectedMasks

[ 20] [RelateObjects] (disabled)

[ 21] [RelateObjects] (disabled)

[ 22] [MeasureObjectIntensity] Measure Intensity of DNA,RNAdye and Membranes under the infected cell masks.

[ 23] [MeasureObjectIntensity] Measure DNA and RNAdye intensity under masks for RNAObjects, MergedRNAobjects and Filtered nuclei objects.

[ 24] [MeasureObjectSizeShape] Measure size and shape of NucleiFiltered and MergedRNAObjects

[ 25] [MeasureObjectIntensityDistribution] Object Intensity desnsity calclulations under infected masks for DNA, RNAdye and BF

[ 26] [MeasureGranularity] Measure Granularity under InfectedMasks for the DNA, RNAdye and BF channels

[ 27] [MeasureTexture] Measure texture under the InfectedCell masks in the DNA, RNAdye and BF channels

[ 28] [ConvertObjectsToImage] (disabled)

[ 29] [ConvertObjectsToImage] (disabled)

[ 30] [ConvertObjectsToImage] (disabled)

[ 31] [SaveImages] (disabled)

[ 32] [SaveImages] (disabled)

[ 33] [SaveImages] (disabled)

[ 34] [ExportToSpreadsheet] Export all data to separate spreadsheets.

[ 35] [ExportToDatabase] Export data and thumbnail images to SQLite database and create props file for use in CellProfiler Analyst.

##Files used and output from this script

This is the neat output from the script I have created and this will always be the most recent version (so may change!!). Older versions are in the old versions directory. Final001.nb.KNOWLESI.html

These are the hex colour codes and how they relate to to the respective clusters Hex.colours.csv

These are the Well definitions as far as I could make out. Introduce various metadata like time. Well\_Defs.csv

This is the list of remaining objects Edgar selected originally. Primarily use to work out how the new clusters relate to them (see table in html file comparing EDmod\_clusters with HC and K clusters). EDmod\_list.csv

Raw data generated by the CellProfiler pipeline and used for this analysis /data/

Frequencies calclulated for clusters per well (HC) /out/KnowlesiHC\_RtsneFreqs2.csv

Static calculated data (saved after t-SNE and clustering 021120). Just used for speed and consistency with graphing. /out/ Knowlesi \_tsne3D\_021120.csv /out/ Knowlesi \_tsne2D\_021120.csv

Output of clusters to image and object numbers. To be imported into CellProfiler Analyst to get images. /out/Knowlesi\_RtsneTrainedSet.csv

Image output from CPA after using the above .csv /out/Knowlsei\_Final001/RGB\_training\_set /out/Knowlsei\_Final001/MGBF\_training\_set
